# Enigmas no longer: using Ultraconserved Elements to place several unusual hawk taxa and address the non-monophyly of the genus *Accipiter* (Accipitriformes: Accipitridae)

**DOI:** 10.1101/2023.07.13.548898

**Authors:** Therese A. Catanach, Matthew R. Halley, Stacy Pirro

## Abstract

Hawks, eagles, and their relatives (Accipitriformes: Accipitridae) are a diverse and charismatic clade of modern birds, with many members that are instantly recognized by the general public. However, surprisingly little is known about the relationships among genera within Accipitridae, and several studies have suggested that some genera (in particular, the megadiverse genus *Accipiter*) are not monophyletic. Here, we combine a new large dataset obtained from Ultraconserved Elements (UCEs), generated from whole genome sequencing (WGS) of 120 species, with publicly available legacy markers (i.e., a suite of commonly sequenced mitochondrial and nuclear genes) to infer a well-supported, time-calibrated phylogeny of 236 extant or recently extinct species. Our densely-sampled phylogeny, which includes 90% of recognized species, confirms the non-monophyly of *Accipiter* and provides a sufficient basis to revise the genus-level taxonomy, such that all genera in Accipitridae represent monophyletic groups.

## INTRODUCTION

Systematists generally agree that genus-rank categories should be composed of monophyletic groups of species, and this is a central tenet of the synapomorphy-based system of classification (Hennig, 1965; Dubois, 1982; Stevens, 1985). However, there is often disagreement about how inclusive such groups should be, and what criteria should be applied to determine generic limits. Some authors have proposed that genera should reflect “adaptive zones” (e.g., Dubois, 1982; Dubois, 1988; Lemen & Freeman, 1984; Miller & Wenzel, 1995), age of divergence (Hennig, 1965; Avise & Johns, 1999), or the overall degree of morphological divergence (e.g., Dubois, 1982). However, each approach has its drawbacks. For example, molecular clocks may produce wildly different estimates for divergence times at a given node, even within the same dataset, depending upon calibration methods (Oatley *et al*., 2015; Mindell *et al*., 2018), and “morphological gaps’’ between monophyletic groups may simply result from incomplete sampling of taxa (e.g., due to extinction). Seeking a practical compromise, Isler *et al*. (2013: 469) encouraged taxonomists to identify generic ranks that “[provide] recognition of phylogenetic relationships, synapomorphic characters, and phenotypic distinctiveness that will best facilitate understanding and communication of relatedness of taxa among analysts, field workers and conservationists.”

We investigated the systematics of a particularly challenging group—the cosmopolitan family Accipitridae (Aves: Accipitriformes), hawks and eagles—which includes multiple large, morphologically diverse genera (e.g., *Accipiter*, *Buteo*), and also a plethora of monotypic genera based on relatively narrow criteria (e.g., *Micronisus*, *Megatriorchis*). The inconsistency of taxonomic practice in Accipitridae has long been a source of taxonomic frustration, as a critique written two centuries ago still rings true:

> “[Authors] are at full liberty to make as many genera or subgenera as they please … But the human mind is ever prone to extremes, and the passion for dividing and subdividing, and giving names, may become as great an evil as that which led the followers of Linnaeus to deprecate *all* division, and to view with abhorrence the slightest attempt to break up the old groups … Fortunately, the only group in Ornithology which has apparently suffered from this evil is that of the *Falconidae* [*sensu lato*, including modern Accipitridae].” (Swainson, 1831: lvii)

Before the mid-20th century, most systematists assumed that falcons (Falconidae) and hawks and eagles (Accipitridae) were sister groups, and thus classified in the order Falconiformes (e.g., Mayr, 1959; but see Starck, 1959). However, based on electrophoretic profiles of egg-white proteins, Sibley (1960) proposed that “Falconiformes [*sensu lato*] may be polyphyletic, the Falconidae possibly being unrelated to the other diurnal birds of prey.” This hypothesis, controversial in its time, has been repeatedly confirmed with molecular phylogenetic analysis (Hackett *et al*., 2008; Jarvis *et al*. 2014; Prum *et al*. 2015); falcons are more closely related to perching birds (Passeriformes) and parrots (Psittaciformes) than hawks. The feather-chewing louse genus *Degeeriella* (Phthiraptera), a parasite of hawks and falcons, is also non-monophyletic; the clade parasitizing falcons is more closely related to the genus *Picicola*, which parasitizes woodpeckers (Catanach & Johnson, 2015). Similarly, New World Vultures (Cathartidae), long thought to be related to storks (Garrod, 1874; Ligon, 1967; Sibley & Ahlquist, 1990; Avise *et al*., 1994), are now understood to be the sister group of Accipitriformes (Jarvis *et al*., 2014; Prum *et al*., 2015).

Modern world checklists including Clements *et al*. (2021), whose nomenclature we follow in this study, currently recognize three families in Accipitriformes: (1) Sagittariidae, a group with unique cranial morphology (Huxley, 1867: 441) consisting of one extant species, the African-endemic Secretarybird *Sagittarius serpentarius* (Miller, 1779), and some extinct taxa from the Oligocene and Miocene (Mourer-Chauviré & Cheneval, 1983); (2) Pandionidae, a group consisting of one extant species, the cosmopolitan Osprey *Pandion haliaetus* (Linnaeus, 1758), and some extinct taxa from the Miocene of North America (Warter, 1976; Becker, 1983); and (3) Accipitridae, a hyperdiverse and globally distributed family that includes hawks, vultures, eagles, and harriers, which contains 249 species in 68 genera. Accipitridae has been divided into several subfamilies and tribes, but these groupings vary widely from study to study. For example Lerner & Mindell (2005) listed 14 subfamilies while Peters (1931) and Mindell *et al*. (2018) recognized only eight subfamilies, the makeup and names of which do not completely match. Mindell *et al*. (2018) also recognized a non-monophyletic group that they referred to as the “transitory Accipitrinae”.

Relative to Sagittariidae and Pandionidae, extant members of Accipitridae exhibit a wide array of morphological, ecological, and behavioral characteristics (del Hoyo *et al*., 1994). Some species have restricted ranges and narrow ecological requirements (e.g., Madagascar Serpent-Eagle, *Eutriorchis astur* Sharpe, 1875, see Kemp & Christie, 2020), while others are ecological generalists with trans-hemispheric distributions (e.g., Golden Eagle *Aquila chrysaetos* (Linnaeus, 1758), see Katzner *et al*., 2020). Most species are solitary predators, with monogamous social and genetic mating systems, but some exhibit cooperative parental care (e.g., Galapagos Hawks *Buteo galapagoensis* (Gould, 1837), see Faaborg *et al*., 1980; DeLay *et al*., 1996), cooperative hunting behavior (e.g., Harris’s Hawk *Parabuteo unicinctus* (Temminck, 1824), see Bednarz, 1988), and even partial frugivory (e.g., Palm-nut Vulture *Gypohierax angolensis* (Gmelin, 1788), see Carneiro *et al*., 2017). Despite this considerable diversity, ornithologists have struggled to resolve relationships within Accipitridae because of high levels of apparent convergence, in external morphology and skeletal structure, likely caused by shared selective pressures that stem inherently from a predatory lifestyle (Holdaway, 1994; Pecsics *et al*., 2019).

Molecular tools have advanced knowledge of some higher level relationships within Accipitridae, but the phylogenetic positions of many taxa remain unresolved. Complete or nearly complete taxon sampling has now been achieved in some genera, but these phylogenies are based on Sanger sequencing of a small number of markers and often lack statistical support for the branching order between genera or groups of genera (e.g., Lerner & Mindell, 2005; Amaral *et al*., 2009). In other bird groups, higher level relationships have been largely resolved by sequencing large numbers of genes, either via genome reduction methods (e.g., Ultraconserved Elements, UCEs) or mining data from whole genome sequencing (Prum *et al*., 2015; Chen *et al*., 2021; Hruska *et al*., 2023). These methods also enable the use of degraded and highly fragmentary samples, like the toepads of study skins (e.g., Catanach *et al*., 2021), whereas Sanger sequencing often requires high quality DNA from fresh or frozen tissue. This is an important breakthrough for studies of Accipitridae systematics because high-quality tissues are lacking for several genera, due to challenges associated with collecting diurnal birds of prey (e.g., heightened legal protection and low population densities). Furthermore, to deal with the scarcity of tissue samples, new methods have been developed to combine “legacy markers” (i.e., Sanger data) with datasets obtained via next generation sequencing (NGS) analyses (Kimball *et al*., 2021).

One group that illustrates these myriad and long standing issues is *Accipiter*, a catch-all genus for (usually) forest-dwelling hawks, into which approximately 50 species have historically been placed (Peters, 1931). For decades, researchers have debated the boundaries of *Accipiter* and its relationship to other genera in Accipitridae, on morphological grounds (Roberts, 1922). Olson (1987) noted that the procoracoid foramen, which is absent in *Accipiter* but present in nearly all other hawk genera, is also absent in some members of *Harpagus* (“kites”) and *Circus* (harriers). However, this was not considered to be strong evidence of a close relationship between *Accipiter* and *Circus*, likely because of their extremely divergent behavior and ecology (Olson, 1987). A subsequent study, focused primarily on osteological characters, concluded that *Accipiter* and *Harpagus* were closely related to the exclusion of *Circus* (Holdaway, 1994). Testing these hypotheses with three molecular markers (ND2, cyt-b, and one nuclear intron), Lerner & Mindell (2005) found strong support for a sister relationship between *Accipiter* and *Circus*, which convinced Olson (2006), but this finding was soon followed by a wave of evidence that *Accipiter*, as traditionally defined (Peters, 1931), does not form a monophyletic group (e.g., Griffiths, 2007; Hugall & Stuart-Fox, 2012; Oatley *et al*., 2015; Mindell *et al*., 2018). As more genes were added to phylogenetic analyses, it became evident that *Circus*, *Megatriorchis*, and *Erythrotriorchis* are nested within a larger “*Accipiter*” clade, which also does not include several other species traditionally placed in *Accipiter*. Furthermore, chewing lice (*Degeeriella*) obtained from Northern Goshawk, *A. gentilis* (Linnaeus, 1758), and Cooper’s Hawk, *A. cooperii* (Bonaparte, 1828), were shown to be more closely related to lice from *Circus* species, than from other “*Accipiter*” species (Catanach & Johnson, 2015).

Despite these findings, previous authors have been hesitant to revise the genus-level taxonomy and nomenclature of Accipitridae for a variety of reasons. Evidence that *Accipiter* is non-monophyletic has primarily come from studies focused on more distantly related clades, which happened to include “*Accipiter*” taxa as outgroups in phylogenetic analyses. In cases where the relationships of *Accipiter* were explicitly addressed, authors have been unwilling to take nomenclatural action because of relatively sparse species-level sampling (e.g., Oatley *et al*., 2015) or uncertainty caused by the exclusion of (or scarcity of available data from) certain enigmatic taxa (Mindell *et al*., 2018). To our knowledge, there have been no attempts to broadly reorganize (under criteria of monophyly) the generic classification of *Accipiter* and related genera. In one case only, have “*Accipiter*” species been reclassified in a new genus after molecular data showed that they were not members of the larger *Accipiter* (*sensu lato*) clade (Hugall & Stuart-Fox, 2012; Oatley, 2015): the osteologically divergent Tiny Hawk, “*Accipiter*” *superciliosus* (Linnaeus, 1766), and Semicollared Hawk, “*Accipiter*” *collaris* (Sclater 1860), are now placed in the genus *Microspizias*, erected by Sangster *et al*. (2021), by some authors.

Here, we estimated phylogenetic relationships among species and genera in Accipitridae by assembling and analyzing multiple molecular datasets: UCEs and legacy markers (nuclear DNA and mitochondrial genomes). Our primary objective was to test the monophyly criterion for each genus-rank taxon, to inform a potential revision of the generic classification of Accipitridae, such that each genus refers to a monophyletic lineage.

## METHODS

### SAMPLE SELECTION

We selected 88 species for whole genome sequencing (WGS) and an additional 8 species for UCE sequencing. All our samples were obtained from vouchered museum specimens, sourced from tissues (69) or study skin toepads (29). We supplemented this dataset by downloading all publicly available raw data (31 samples) from the European Nucleotide Archive (https://www.ebi.ac.uk/ena/browser/), resulting in a combined dataset (supplemental table 1) that included samples from 41 genera in Accipitridae (ingroup) and 7 genera in Pandionidae, Sagittariidae, and Cathartidae (outgroup). When multiple genomes were available for the same species, we selected the sample with most UCEs assembled (i.e., for the UCE analysis), but included all available samples for gene tree analyses involving legacy markers.

To include species for which WGS sequencing had not yet been performed, we downloaded all publicly available sequence data (i.e., “legacy markers”) from Accipitriformes and Cathartiformes from Genbank (https://www.ncbi.nlm.nih.gov/genbank/), with a submission cutoff date of 28 February 2022. We also downloaded sequence data from BOLD (http://www.boldsystems.org/), if they were obtained from a taxon not represented in the Genbank dataset. We included all available mitochondrial data (with the exception of control regions and tRNAs), but only five nuclear genes (i.e., those for which broad sampling was available): portions of beta-fibrinogen (FGB) 4-8 (exons and introns), myoglobin (MB) exon 2 and intron, myelocytomatosis viral oncogene-like protein (c-myc), transforming growth factor beta 2 (TGFb2) intron-5, and recombination activating protein 1 (RAG1).

### EXTRACTION, SEQUENCING, AND LIBRARY PROCESSING

We extracted DNA from all samples with Qiagen DNAeasy Kits (Germantown, MD) by following the manufacturer’s instructions, with the exception of toepad samples used for UCE sequencing, which we extracted using a phenol-chloroform protocol followed by bead cleanup (Tsai *et al*., 2019) or a modified QIAamp DNA Micro Kit (Germantown, MD) developed by Andres Cuervo (personal communication). For WGS sequencing, Illumina libraries were constructed with the Illumina TruSeq kit with standard adapters, and sequencing was performed on the Illumina X-Ten platform, at Genewiz (South Plainfield, NJ, USA). For UCE samples, library preparation and 150 bp paired-end reads on an Illumina HiSeq 3000/4000 was performed by Rapid Genomics (Gainesville, FL) using either the 2.5k tetrapod kit (which also includes ∼100 Avian exons) or the 5k tetrapod probe set (Faircloth *et al*., 2012). We first removed duplicate reads using the scriptfastqSplitDups.py from mcscript (https://github.com/McIntyre-Lab/mcscript), then used bbduk.sh from bbmap (Bushnell, 2014) to remove adaptors. The only exceptions were some extremely large (>150 gb) libraries, which we processed without duplicate removal after finding (via fastqc) that they had low levels of read duplication (Andrews, 2010).

### UCE ASSEMBLY

We assembled UCEs using aTRAM2 (Allen *et al*., 2018). For our target sequences, we selected hawk-specific versions of the probes in the 5k tetrapod probe set. We used the Phyluce pipeline (Faircloth, 2016) to assemble UCEs from several high-quality hawk tissue samples, and then selected the longest representative of each UCE to serve as a target sequence for UCE assembly for other species in the study (Catanach *et al*., 2021). Using these targets, we then performed five iterations of BLAST queries against each shard (created by partitioning a large dataset into several smaller datasets to ease computational requirements). Each shard was approximately 125 MB and each BLAST query was capped at 4,000 sequences per shard to decrease runtime. We then created UCE alignments with a set of custom scripts (https://github.com/juliema/phylogenomics_scripts), which extracted the longest assembly from each UCE, combined them across samples, and aligned them using MAFFT (Katoh & Standley, 2013).

We checked each UCE alignment by eye and removed unalignable portions of individual sequences. In a few cases, we encountered sets of assembled UCE sequences that matched each other but differed drastically from other sequences. We assumed that this was an artifact of assembly, and not biological signal, because UCEs are conserved across all tetrapods (Faircloth *et al*., 2012) and samples from the same avian order (Accipitriformes) are not expected to exhibit extremely different motifs and these UCEs were removed. We then used the Python 3 package Fastaq (https://github.com/sanger-pathogens/Fastaq) to create individual UCE alignments with the selected samples. Within each alignment, individual UCE sequences varied widely in length (from ∼100 bp to >6,000 bp). Therefore, to limit the amount of missing data, we used trimAL (Capella-Gutiérrez *et al*., 2009) to remove columns with more than 30% missing data and excluded all UCEs that did not include at least 119 of the 120 taxa in the dataset. The final dataset consisted of 1,582 UCEs and a total of 1,752,961 columns.

### LEGACY SEQUENCE ASSEMBLY FROM NGS SAMPLES

We used readmapping in Geneious 8.1.9 (Biomatters Limited, Auckland, NZ) to assemble mitochondrial genomes (mitogenomes) and legacy markers. We used a set of reference mitogenomes and legacy gene sequences from Genbank, set the sensitivity to allow no more than 10% of bases to be mismatches, and ran five iterations of the readmapping function. During this process, virtually all of the legacy markers assembled to completion but, in a few instances, small portions of the mitogenome (excluding control region) were not assembled. For these samples, we repeated the readmapping step with the assembled portion of the incomplete gene as the reference, to ensure no viable data were missed. After the assembly was complete, we annotated the genomes by first creating consensus sequences, then using the “apply annotation” function, the source being the same reference mitogenome used for the assembly. Finally, we aligned our genome and the reference genome to each other and visually checked the annotation (i.e., by eye). Due to genome rearrangements and difficulty aligning the control regions, we did not attempt to analyze the mitogenome as a whole. Rather, we extracted each gene and ribosomal RNA, placed them in a file combining all samples, and aligned them using MAFFT.

### INTEGRATION OF LEGACY AND NGS DATASETS

Before integrating the data obtained via readmapping legacy genes to our WGS and UCE libraries, we performed several quality control steps. First, we updated the taxonomy to match Clements *et al*. (2021) and, where possible, assigned individual samples to subspecies. This step was necessary because some currently recognized species in Accipitridae are evidently not monophyletic (e.g., Kunz *et al*. 2019). To accomplish this, we extracted the collection locality and date directly from Genbank, associated publication(s), and/or museum databases using the provided voucher information. If subspecies could not be conclusively determined, we assigned the sample to species only. When available, notated each sample in the alignment with its voucher number or other individual identification information.

Next, we added legacy markers assembled from NGS samples by using the read-mapping process (see above) and aligned individual genes using MAFFT. Then, for quality control, we applied a single model (GTR+I+G, without model testing) to estimate gene trees using IQ-TREE. This enabled questionable sequences to be quickly identified and addressed. First, after this initial quality control analysis, we removed all samples on extremely long branches, which likely resulted from contamination or poor sequence quality. Next, we investigated all instances where an individual sample was not placed with other samples thought to be its closest relatives. Often, these errors turned out to be misidentifications, in which case we updated the identification in our dataset. If a problematic sequence was sourced from a specimen that had been sequenced for multiple genes (based on voucher information), we examined its placement in the other gene trees to potentially shed light on its identity, especially when no other samples of that species were available for that particular gene. If we were unable to identify the cause of the error, despite these efforts, we removed the samples from the dataset. We continued this process until no long branches remained and we were confident that all misidentifications were removed. Lastly, we excluded extremely short samples (< 75 bases long).

Next, after the quality control process was complete, we used Seaview v.4 (Gouy *et al*., 2010) to concatenate the legacy data by voucher (all genes sequenced from a given individual were concatenated). This produced a single alignment sourced from (potentially) over 5,000 individual hawks and vultures. Not all Genbank records contained voucher numbers, and we only assumed that sequences came from the same individual when specimen voucher information was explicitly stated, or samples were identified as coming from an individual with a unique code such as a banding number so this is likely an overestimate. As expected in supermatrix approaches, the majority of our alignment was missing data (85.72% in the mitochondrial portion, and even more when nuclear data was included) and most samples were represented by a single gene. Nevertheless, a majority of species were represented by several samples with differing sets of sequenced genes. Therefore, when necessary, we combined samples from multiple individuals of the same species, so that each species was represented by a single (sometimes composite/chimerical) sequence that included as much data as possible for that taxon (Fig. 1).

**Figure 1.**
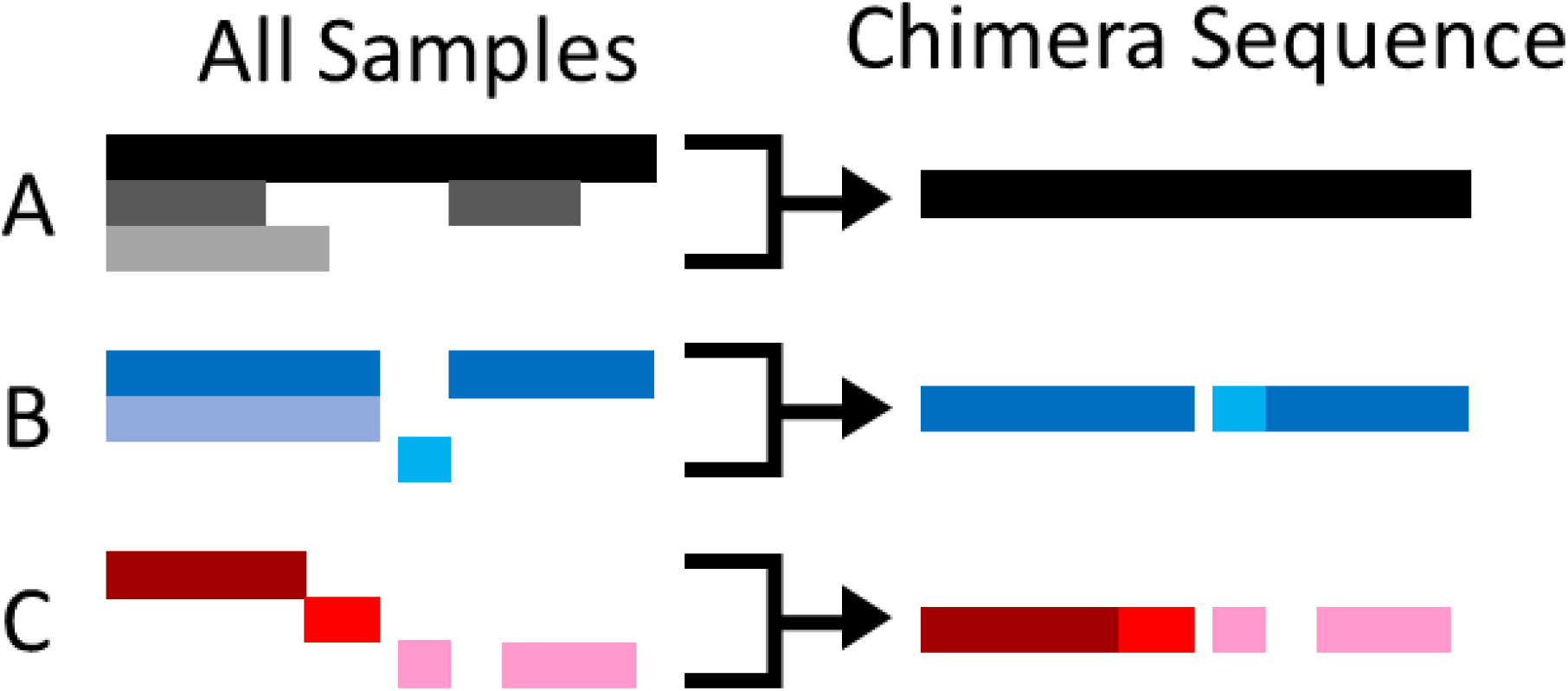
Schematic showing three methods for assembling composite (chimera) sequences. In each case (A–C), samples from different individuals of the same species (denoted by different shades of the same color) were combined to create a single composite sequence containing the most complete set of sequence data possible In scenario A, a full mitochondrial genome was available (black bar) and therefore chosen to represent the species in the final alignment (i.e., no composite was necessary). In scenario B, because the middle sample was of identical length to the corresponding fragment in the top sample, which had additional fragments available, the top sample was retained in the composite sequence; the lowest sample did not overlap with any sequenced portions of the top sample, and was therefore also retained in the composite. In scenario C, the three sequenced individuals did not share any overlapping sequenced regions, so all samples were combined to form the chimeric sequence.

For species represented in our NGS dataset, we preferentially used legacy data assembled from the same NGS sample, with one exception. For *Accipiter bicolor* (Vieillot, 1817), we used data obtained from a different NGS sample (LSUMZ 24224), from which we assembled fewer UCEs but more legacy markers compared to FMNH 260973 in our NGS dataset. In a few cases, the NGS sample had unsequenced regions and, when possible, we filled these areas with sequence data from Genbank samples. It is important to note that, when gene tree analysis revealed that a species was not monophyletic, we created a composite sequence from only a monophyletic lineage (e.g., from a single subspecies or group of subspecies that form a clade). The resulting alignment (hereafter called the “236 species alignment”) contained 36.74 and 39.31% missing data in the mitochondrial and nuclear portions, respectively. Supplemental Table 1 lists all accession numbers for samples included in this final alignment.

### PHYLOGENETIC INFERENCE

To estimate a UCE phylogeny, we concatenated the 1,494 UCE alignments into a single partitioned file using catfasta2phyml (https://github.com/nylander/catfasta2phyml). We used modelfinder (Kalyaanamoorthy *et al*., 2017) as implemented in IQ-TREE 1.6 (Nguyen, 2015) to select the best substitution model for each UCE, followed by tree inference in IQ-TREE 1.6 using CIPRES (Miller *et al*., 2010). We also performed 1,000 ultrafast bootstraps (Hoang *et al*., 2018) to assess statistical support for each node. Hereafter, we refer to this phylogeny as the “120 species UCE phylogeny”.

While legacy data may be useful for determining relationships among recently diverged groups, they have been insufficient to resolve deeper (older) relationships between clades, in most phylogenies of Accipitriformes published to date (e.g., Lerner & Mindell, 2005; Starikov & Wink, 2020). Our legacy dataset was no exception. Preliminary analysis of commonly used nuclear and mitochondrial genes, using IQ-TREE 1.6 (Nguyen, 2015) for model selection and tree inference, produced phylogenies with no statistical support for many key nodes. These topologies were also impacted by the nonrandom distribution of missing data. Therefore, to determine the placement of the 116 species represented only by legacy markers, relative to the well-supported 120 species UCE phylogeny, we employed the UCE tree as a “backbone” by using the –tree-constraint command in RAxML-NG v1.1.0 (Kozlov *et al*., 2019) as implemented on CIPRES (Miller *et al*., 2010). We then selected a GTR+G model for each legacy gene and performed a (constrained) phylogenetic analysis on the 236 species alignment, which produced a backbone phylogeny that matched the topology inferred using UCEs, while allowing the legacy data to determine the placement of the taxa lacking NGS data.

In our first attempt, three species were unexpectedly placed in the resulting phylogeny (i.e., placed in clades not found in any gene tree). One species, *Accipiter madagascariensis* Verreaux, 1833, was represented by a miniscule amount of data (298 bp of CO1). Similarly, *A*. *poliogaster* (Temminck, 1824) and *A*. *ovampensis* Gurney, 1875 were represented by a small fragment of CO1 (298–652 bp) and a fragment of myc (1074 bp). In the CO1 gene tree, this species were clustered unambiguously in a clade that also included *A. nisus* (Linnaeus, 1758), *A. rufiventris* Smith, 1830, and *A. striatus* Vieillot, 1808; whereas, in the myc gene tree, *A*. *ovampensis* was included in the *A*. *nisus* clade and the placement of *A*. *poliogaster* was equivocal (although within *Accipiter senus lato*). Therefore, we reanalyzed the 236 species alignment with an additional topological constraint, requiring *A. madagascariensis*, *A. ovampensis*, and *A. poliogaster* to be placed within the clade containing *A*. *nisus*, *A. rufiventris*, and *A. striatus,* but did not assign them to any particular position within that clade. Finally, because we used multiple topological constraints, we did not calculate support values for nodes in the resulting phylogeny.

### DIVERGENCE TIME ESTIMATION

We performed divergence time estimation on the 236 species phylogeny with BEAST 1.7.5, by restricting our dataset to the most commonly sequenced genes (COI, ND1, ND2, ND6, cytb, RAG1, and myc) and holding the topology constant. For each gene, we unlinked substitution rates and clock rates, assigned a GTR+G substitution model, and estimated divergence times with a strict clock. We calibrated the molecular clock with two fossils: (1) *Circaetus rhodopensis* (Boev, 2012), which is informative of the split between the *Circaetus* + *Dryotriorchis* clade and *Terathopius ecaudatus* (Daudin, 1800), implemented using a lognormal prior with a mean of 7.5 MYA, standard deviation of 0.25, and an offset of 7.25; (2) *Aegypius varswaterensis* (Manegold *et al*., 2014), which is informative of the split between *Aegypius* and *Torgos*, was implemented using a lognormal prior with a mean of 5 MYA, standard deviation of 0.25, and an offset of 3.6. We also placed a normal prior on the root of the tree with a mean of 60.34 MYA, and a standard deviation of 1.61 (Knapp *et al*., 2019). We then performed 10 million MCMC steps, sampling every 1,000 generations. We used TreeAnnotator to annotate the maximum clade credibility tree after discarding 25% of the trees as burn-in.

## RESULTS

### PHYLOGENETIC HYPOTHESES

Our 120 species UCE phylogeny (Figure 2; Supplemental Figure 1 for detailed divergence estimates) was well supported (bootstrap values = 100) at virtually every node. Only five nodes had less than perfect support, and four of those represented splits between closely related species or groups of species (two within *Buteo*, one within *Buteogallus*, one in *Gyps*). The fifth node with less than perfect support represented the split between two closely related “kites” (*Helicolestes hamatus* (Temminck, 1821), *Rostrhamus sociabilis* (Vieillot, 1817)). Of the 19 genera for which multiple species were sampled in our 120 species UCE phylogeny, all but *Accipiter* were monophyletic. The non-monophyly of *Accipiter* was primarily caused by the embedded placement of *Circus, Erythrotriorchis*, and *Megatriorchis* (see below). Furthermore, in the 236 species phylogeny, the genus *Circaetus* was not monophyletic, because *Dryotriorchus* was embedded within it.

**Figure 2.**
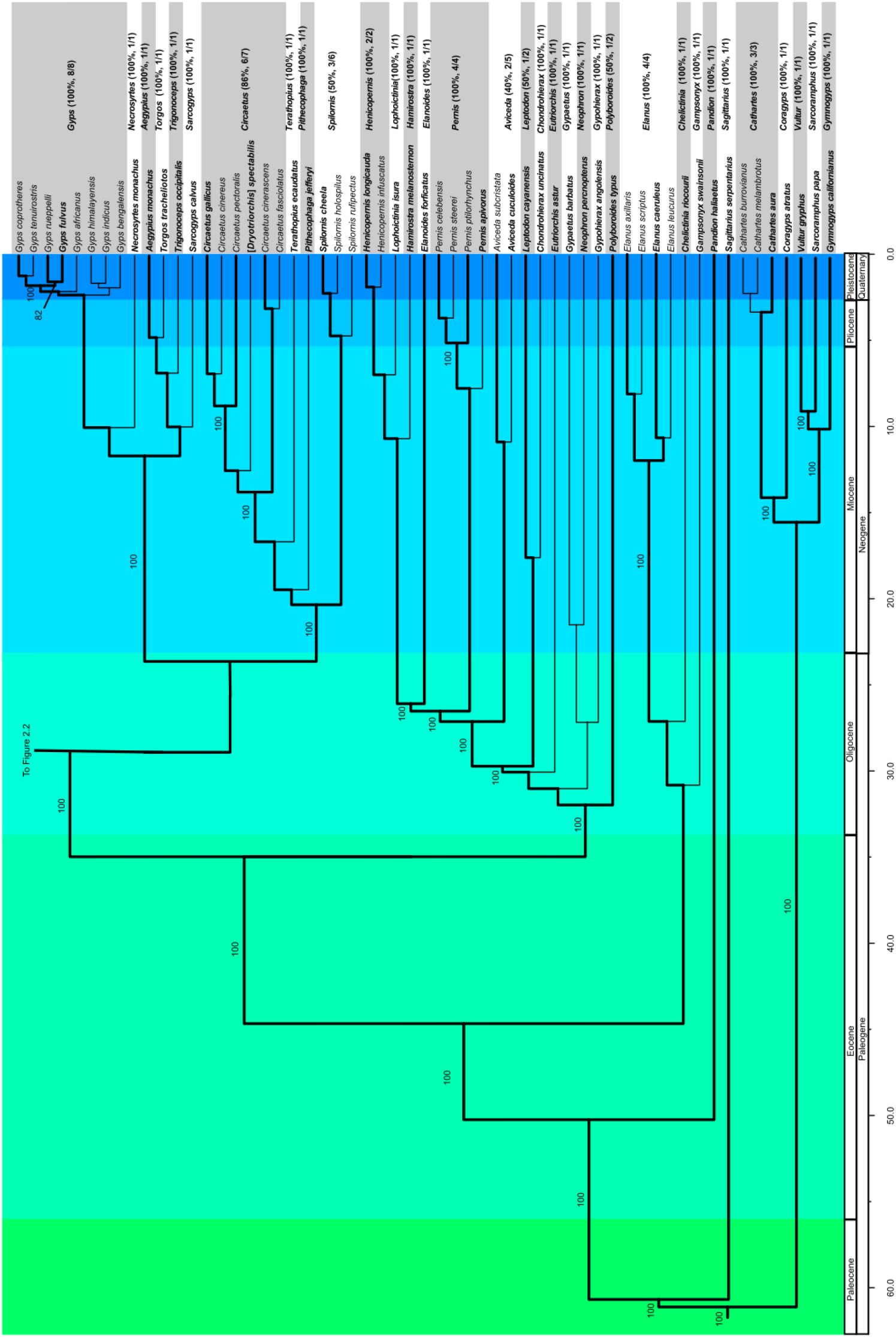
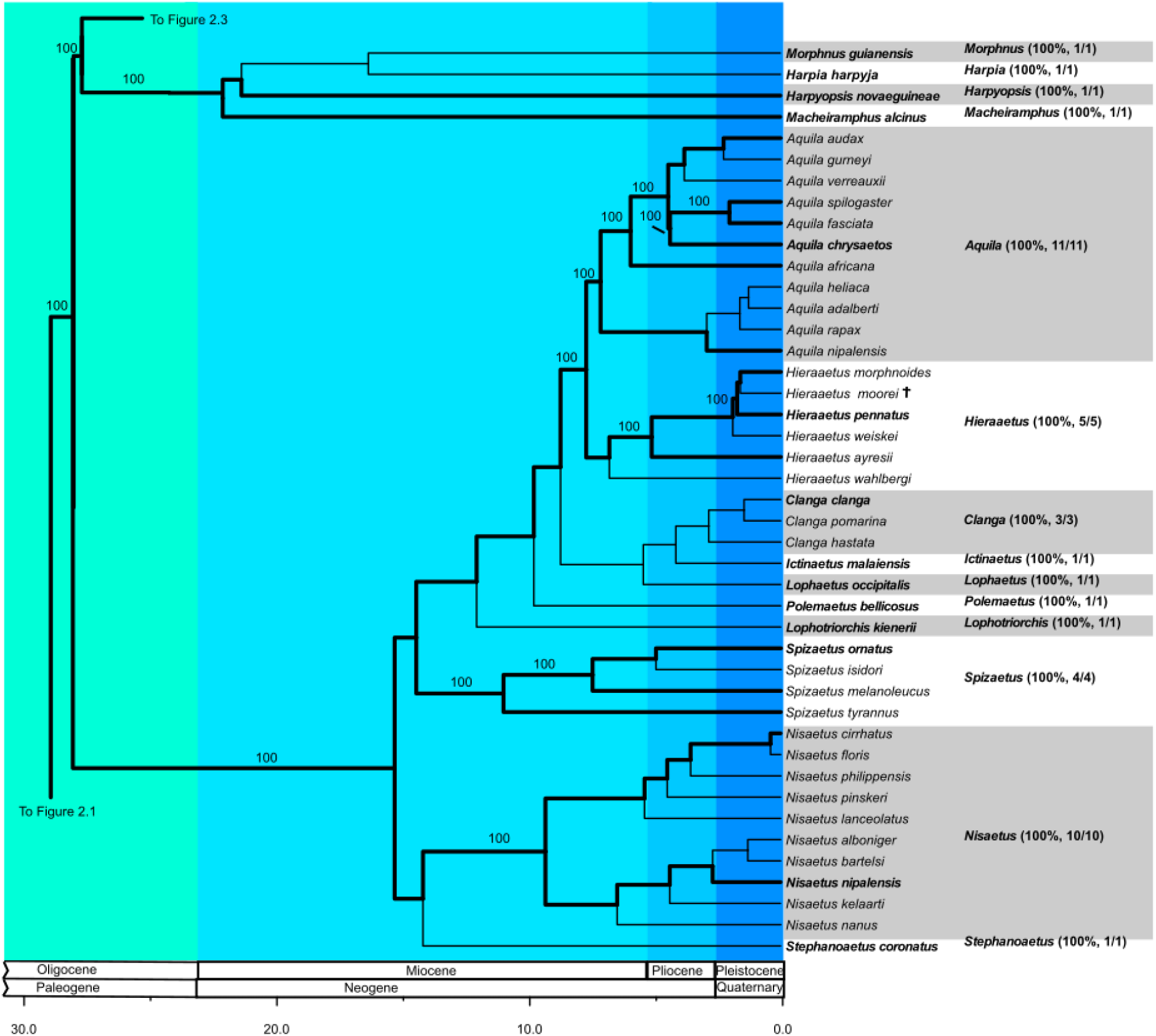
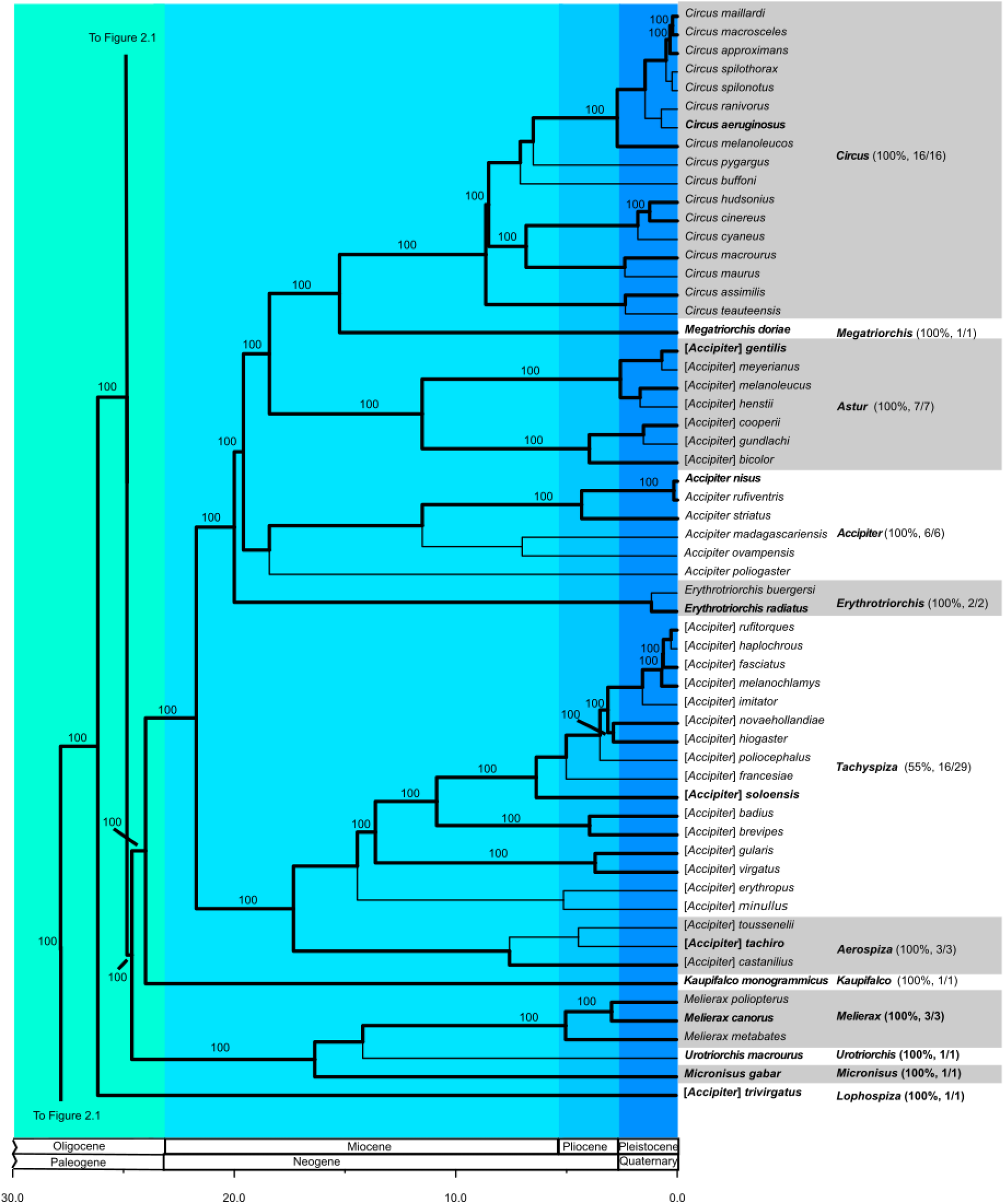
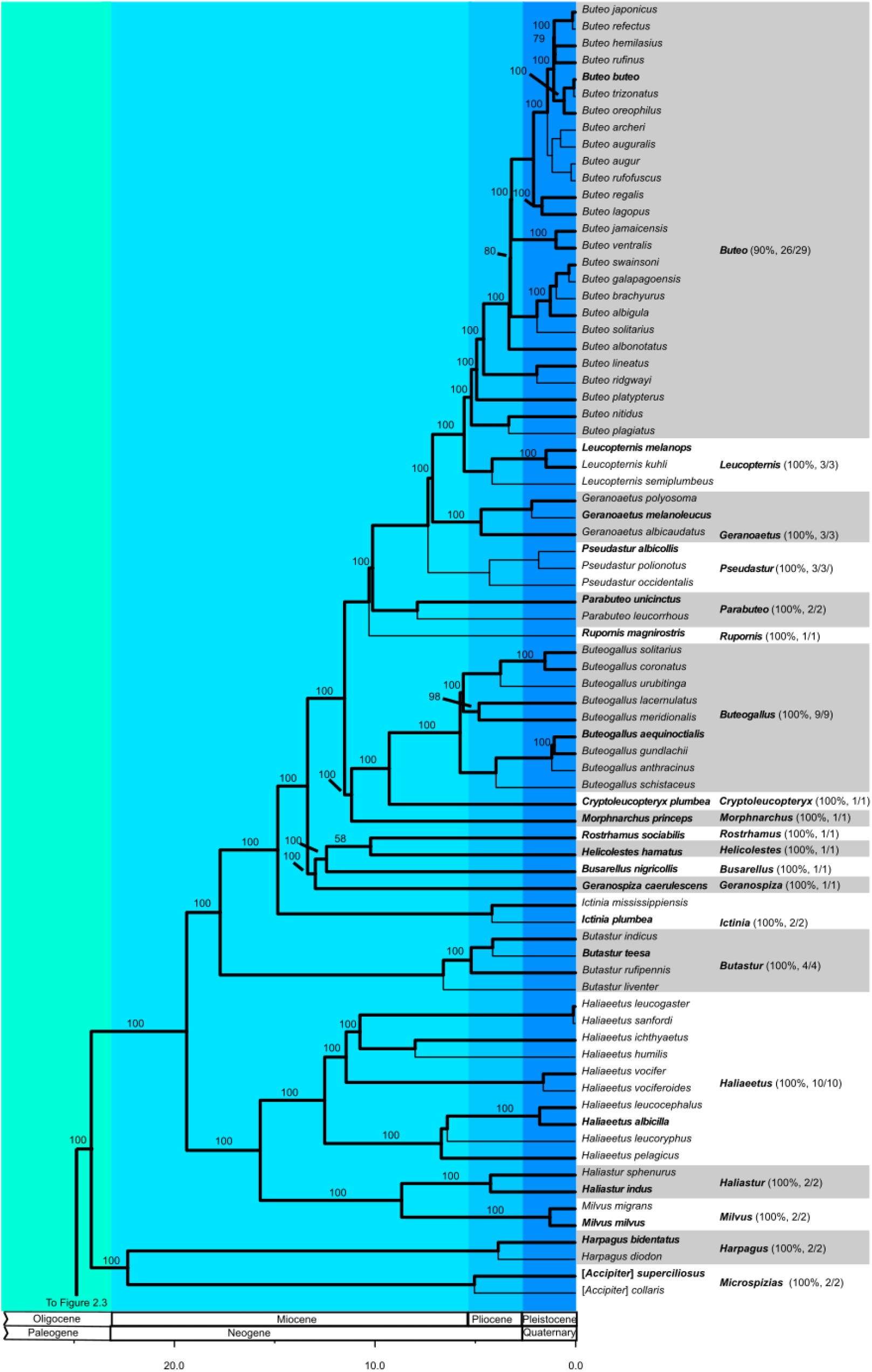
Phylogeny of the Accipitridae with outgroup sequences of Pandionidae, Sagittariidae, and Cathartiformes. Thickened lines show the topology of the 120 species UCE phylogeny. Thinner lines show the placement of taxa represented only by legacy markers (see Methods). UCE bootstrap values are shown above each node. Support values were not calculated for nodes connecting taxa represented only by legacy data. Colors represent geologic epochs. Gray and white bars denote monophyletic groups that are classified at the rank of genus in our taxonomic revision. For each group, the genus name that holds taxonomic priority is used, and the type species is shown in boldface.

The order Cathartiformes formed a clade that was sister to Accipitriformes. Within Accipitriformes, the family Sagitariidae was sister to the rest of Accipitriformes, and Pandionidae was sister to the rest of Accipitridae. Several large clades within Accipitridae corresponded roughly (except as noted) to the subfamilies recognized by Mindell *et al*. (2018) and Lerner & Mindell (2005). Hereafter, for convenience, genera in boldface denote those included in the 120 species UCE phylogeny.

The most basal split within Accipitridae separated the monophyletic subfamily Elaninae (composed of ***Elanus****, Gampsonyx, and Chelictinia*) from the rest of the taxa. Within the sister group of Elaninae, a clade containing all genera traditionally placed in Gypaetinae (***Polyboroides***, *Gypohierax*, *Neophron*, *Gypaetus*) and Perninae (*Eutriorchis*, ***Leptodon***, *Chondrohierax*, ***Elanoides***, ***Pernis***, ***Aviceda***, ***Hamirostra***, *Lophoictinia*, ***Henicopernis***) formed the sister group of the remainder of Accipitridae. However, *Polyboroides typus* Smith, 1829 was placed as sister to the rest of the Gypaetinae + Perninae clade, which rendered Gypaetinae non-monophyletic.

Within the sister group of Elaninae + Gypaetinae + Perninae, a clade containing all members of the subfamilies Circaetinae (***Spilornis***, *Pithecophaga*, *Terathopius*, ***Circaetus***, *Dryotriorchis*) and Aegypiinae (*Sarcogyps*, *Trigonoceps*, *Torgos*, ***Aegypius***, *Necrosyrtes*, ***Gyps***), which were reciprocally monophyletic, formed the sister group to the remainder of Accipitridae. *Dryotriorchis spectabilis* (Schlegel, 1863) was embedded within the genus ***Circaetus***, rendering it paraphyletic. Within the sister group of Elaninae + Gypaetinae + Perninae + Circaetinae + Aegypiinae, a clade containing the subfamily Aquilinae (*Stephanoaetus*, ***Nisaetus***, ***Spizaetus***, *Lophotriorchis*, *Polemaetus*, *Lophaetus*, *Ictinaetus*, *Clanga*, ***Hieraaetus***, ***Aquila***) formed the sister group to the remainder of Accipitridae.

Within the sister group of Elaninae + Gypaetinae + Perninae + Circaetinae + Aegypiinae + Aquilinae, a clade composed of genera traditionally placed in the subfamily Harpiinae (***Macheiramphus***, ***Harpyopsis***, *Morphnus*, *Harpia*) was the sister group of the remainder of Accipitridae. Within the sister group of Elaninae + Gypaetinae + Perninae + Circaetinae + Aegypiinae + Aquilinae + Harpiinae, the monotypic “*Accipiter*” *trivirgatus* was recovered as the sister group to the remainder of Accipitridae, which were divided into three major clades. The first of these was the “*Accipiter*” complex (***Kaupifalco***, ***Micronisus***, ***Melierax***, *Urotriorchis*, ***Erythrotriorchis***, ***Megatriorchis***, ***Circus***, ***Accipiter***), which included taxa traditionally placed in three subfamilies (Melieraxinae, Circinae, and Accipitrinae).

The final two major clades were reciprocally monophyletic. One contained the genera ***Harpagus*** and two “*Accipiter*” species that were recently reclassified in the genus ***Microspizias*** (see Sangster *et al*., 2021): [*A.*] *superciliosus* and [*A.*] *collaris*, while the final clade contains all Buteoninae, which has occasionally been divided into two tribes, Milvini (***Milvus***, ***Haliastur***, ***Haliaeetus***) and Buteonini (***Butastur***, ***Ictinia***, ***Busarellus***, ***Rostrhamus***, ***Helicolestes***, ***Geranospiza***, ***Cryptoleucopteryx***, ***Buteogallus***, ***Morphnarchus***, *Rupornis*, ***Parabuteo***, ***Geranoaetus***, *Pseudastur*, ***Leucopternis***, ***Buteo***), both of which were recovered as monophyletic in our dataset.

### DIVERGENCE TIMING

The orders Cathartiformes and Accipitriformes were estimated to have diverged at 61.1 MYA. The common ancestor of extant taxa within Cathartidae was estimated at 15.6 MYA. The family *Sagittariidae* diverged from the rest of Accipitriformes at 60.7 MYA, and *Pandionidae* diverged from Accipitridae at 50.2 MYA. The clades corresponding to subfamilies within Accipitridae diverged between 23.7 and 32 MYA, with the exception of Elaninae which split from the rest of Accipitridae at 44.7 MYA. The majority of generic level splits occurred at least 5 MYA, although in some cases several million years passed between the inferred origin of the genus (i.e., the node uniting it with its sister group) and the most recent common ancestor of its extant (or recently extinct) members.

## DISCUSSION

Our phylogeny was topologically similar to previously published studies of Accipitridae and included the non-monophyly of *Accipiter*. However, by including several enigmatic species, we removed a primary source of uncertainty that prevented former authors from reconciling the genus-level nomenclature with phylogenetic data. Our dense sampling framework included 90% of extant Accipitridae species (225/249, following Clements *et al*., 2021), plus two extinct species, all of which were represented by at least one gene. Approximately half of these species were included in our UCE dataset, which enabled us to test the monophyly criterion with greater confidence. These data are sufficient to recommend a conservative revision of the generic classification, which divides the non-monophyletic *Accipiter* into multiple genera that reflect evolutionary relationships. Hereafter, we discuss our results and taxonomic proposals within the context of each major clade.

### CATHARTIFORMES

Our analysis, which included UCE data from all five extant genera and legacy data from all currently recognized extant species, recovered Cathartiformes as the sister clade to Accipitriformes. This finding is supported by multiple morphological synapomorphies (e.g., Griffiths 1994). Within Cathartiformes, we found that Black Vulture, *Coragyps atratus* (Bechstein, 1793), was sister to *Cathartes*, and these two genera were sister to a clade containing the remaining New World Vultures (i.e., *Gymnogyps* + *Vultur* + *Sarcoramphus*) — a hypothesis previously proposed by Johnson *et al*. (2016). We also found strong evidence (Bootstrap Value = 100) that *Vultur* was sister to *Gymnogyps* + *Sarcoramphus*, corroborating the poorly supported topology recovered by Johnson *et al*. (2016). Our estimated divergence time for the split between the two main clades of Cathartiformes (15.6 MYA) was similar to that of Johnson *et al*. (2016), but they estimated a older date for the Cathartiformes and Accipitriformes split (∼69 MYA), whereas our estimate (61.1 MYA) was more similar to those of Jarvis *et al*. (2014) and Knapp *et al*. (2019).

### SAGITARIIDAE & PANDIONIDAE

The phylogenetic placement of these monotypic families in our phylogeny was identical to previous studies. We estimated that Sagittariidae diverged from the Pandionidae + Accipitridae clade around 60.7 MYA, shortly after the split between Accipitriformes and Cathartiformes; and we estimated that Pandionidae diverged from Accipitridae around 50.2 MYA. These estimates were similar to those inferred by Johnson *et al*. (2016) and Knapp *et al*. (2019), but considerably older than those of Mindell *et al*. (2018) (*c.*45 MYA and *c.*38 MYA, respectively) and Prum *et al*. (2015) (*c.*40 MYA and *c.*28 MYA, respectively). These discrepancies probably resulted from different calibration methods. In our study, following Johnson *et al*. (2016) and Knapp *et al*. (2019), we placed a prior on the split between Cathartiformes and Accipitriformes, whereas Mindell *et al*. (2018) and Prum *et al*. (2015) did not. Johnson *et al*. (2016) also used several fossil calibrations to inform the divergence estimates. Interestingly, although we used different priors and fossils than Johnson *et al*. (2016), we arrived at nearly identical divergence estimates for these two splits.

### ELANINAE

Within the Elaninae, the monotypic Scissor-tailed Kite, *Chelictinia riocourii* (Temminck, 1821), was sister to the monophyletic genus *Elanus*, which is comprised of four species, and Pearl Kite, *Gampsonyx swainsonii* Vigors, 1825, was sister to the *Chelictinia* + *Elanus* clade. This identical arrangement was proposed by Starikov & Wink (2020), based on analysis of one nuclear and two mitochondrial genes, although they lacked data from Letter-winged Kite, *E*. *scriptus* Gould, 1842. Our results were also similar to Mindell *et al*. (2018), who lacked data from *C. riocourii* and Black-shouldered Kite, *E*. *axillaris* (Latham, 1802). Divergence estimates published by these authors differed from each other and our own. Our estimate for the divergence of Elaninae from the rest of Accipitridae (44.7 MYA) was roughly twice as old as that of Starikov & Wink (2020), and the estimate of Mindell *et al*. (2018) was intermediate. Conversely, our estimate for the divergence of *Elanus* and *Chelictinia* was about twice as old as the estimate of Mindell *et al*. (2018), whereas Starikov & Wink (2020) provided an intermediate estimate. These discrepancies are not surprising, as each study used different calibration methods.

### GYPAETINAE, PERNINAE, & POLYBOROIDINAE

Within the clade containing most species typically placed in the subfamilies Gypaetinae, Perninae, and Polyboroidinae (e.g., Brown & Amadon, 1968; Lerner & Mindell, 2005), we found that African Harrier-Hawk (*Polyboroides typus*) was sister to a clade containing the rest of the species. This relationship was recovered by Lerner & Mindell (2005) using three legacy markers, although Mindell *et al*. (2018) later found evidence that *P. typus* was sister to Palm-nut Vulture (*Gypohierax angolensis*), supporting a monophyletic Gypaetinae (i.e., *Polyboroides* + *Gypohierax* + *Neophron* + *Gypaetus*). Our results only support a monophyletic Gypaetinae if *Polyboroides* is excluded, although we think additional sampling is needed to be confident about this result. Three members of the “true” Gypaetinae were represented in our analysis by six or more legacy genes, but we lacked UCE data for this group and therefore hesitate to recognize a monotypic subfamily for *P. typus* (Polyboroidinae) at this time, as some authors have done (e.g., Brown & Amadon, 1968). Nevertheless, we acknowledge that these three monotypic genera are highly divergent from each other, morphologically and ecologically (e.g., van Lawick-Goodall & van Lawick, 1966; Stoyanova *et al*., 2010). Within the “Gypaetinae” clade (i.e., minus *Polyboroides*), we recovered a sister relationship between Bearded Vulture, *Gypaetus barbatus* (Linnaeus, 1758), and Egyptian Vulture, *Neophron percnopterus* (Linnaeus, 1758), and found that this clade was sister to Palm-nut Vulture (*Gypohierax angolensis*).

We did not find support for the inclusion of Madagascar Serpent-Eagle, *Eutriorchis astur*, in the Circaetinae (*contra* Brown & Amadon, 1968), nor in the Gypaetinae (*contra* Lerner & Mindell, 2005), nor nested within the Perninae, as sister to a clade containing Grey-headed Kite, *Leptodon cayanensis* (Latham, 1790), and Hook-billed Kite, *Chondrohierax uncinatus* (Temminck, 1822), as proposed by Mindell *et al*. (2018). Rather, we recovered *E. astur* as the sister group of a monophyletic Perninae containing the genera *Henicopernis*, *Lophoictinia*, *Hamirostra*, *Elanoides*, *Pernis*, *Aviceda*, *Leptodon*, and *Chondrohierax*. Because it was only represented by legacy data, we are unable to conclusively resolve the placement of *E. astur* at this time. Within the remaining clade of Perninae (i.e., sister to *E. astur* in our phylogeny), we found similar (but not identical) relationships among species to Lerner & Mindell (2005) and Mindell *et al*. (2018), albeit with denser taxon sampling. One notable difference was our placement of Swallow-tailed Kite, *Elanoides forficatus* (Linnaeus, 1758), as sister to a clade containing *Hamirostra*, *Lophoictinia*, and *Henicopernis*, whereas Mindell *et al*. (2018) found *E. forficatus* to be sister to the rest of Perninae, and Lerner & Mindell (2005) did not include *Henicopernis*. Furthermore, we found that Pacific Baza, *Aviceda subcristata* (Gould, 1838), and African Cuckoo-Hawk, *A. cuculoides* Swainson, 1837, were sisters (i.e., the genus *Aviceda* was monophyletic), whereas Mindell *et al*. (2018) placed *A. subcristata* as sister to the *Henicopernis* + *Lophoictinia* + *Hamirostra* clade, and Lerner & Mindell (2005) did not include any *Aviceda* samples in their analysis. This situation highlights the potential erroneous topologies that may arise in phylogenetic studies that rely on a sparsely populated supermatrix; notably, in the Mindell *et al*. (2018) dataset there was no gene overlap between the two included *Aviceda* species. Our approach of mining legacy markers from NGS data allowed us to fill in these gaps, resulting in a phylogeny that confidently placed these two species as sister groups.

### CIRCAETINAE

Within a clade containing most of the genera formerly placed in the subfamily Circaetinae (excluding *Eutriorchis*, *contra* Lerner & Mindell, 2005), samples from three species in the genus *Spilornis*, which is composed primarily of southeast Asian island endemics, formed a clade that was sister to a clade containing samples from the genera *Pithecophaga*, *Terathopius*, *Circaetus*, and *Dryotriorchis*. A sample of Congo Serpent Eagle (*Dryotriorchis spectabilis*), which had been classified in *Circaetus* prior to Mindell *et al*. (2018), was nested within a clade of *Circaetus* samples. A sample of Bateleur, *Terathopius ecaudatus*, was sister to *Circaetus* (*sensu lato*, including [*D.*] *spectabilis*), and a sample of Philippine Eagle, *Pithecophaga jefferyi* Ogilvie-Grant, 1896, was sister to the *Circaetus* + *Terathopius* clade. These results agreed with previous phylogenies in the placement of *Pithecophaga* as sister to a clade containing *Terathopius* and *Circaetus* (Lerner & Mindell, 2005; Mindell *et al.,* 2018), but not with respect to the arrangement of the other species. Mindell & Lerner (2005) did not sample [*D.*] *spectabilis*, whereas Mindell *et al*. (2018) found that it was sister to the *Terathopius* + *Circaetus* clade. Although [*D.*] *spectabilis* was represented only by legacy markers in our study, we have UCE data from several *Circaetus* species, including from both clades created by the inclusion of [*D.*] *spectabilis*. Therefore, we think it is appropriate to restore [*D.*] *spectabilis* to *Circaetus*, according to tradition. Our divergence estimate for the crown age of Circaetinae was 20.6 MYA, which is substantially older than Mindell *et al*.’s (2018) estimated date of *c.*14 MYA. We used the same fossil calibration as Mindell *et al*. (2018), for the split between *Circaetus* + *Terathopius*, but our tree topologies were in conflict for this node.

### AEGYPIINAE

A clade containing all six genera traditionally placed in the subfamily Aegypilnae (*Trigonoceps, Gyps*, *Necrosyrtes*, *Aegypius*, *Torgos*, *Sarcogyps*) contained two subclades, the first containing the (reciprocally monophyletic) sibling genera *Gyps* and *Necrosyrtes*, and the other containing the genera *Aegypius*, *Torgos*, *Trigonoceps*, and *Sarcogyps*, arranged in a nested pattern. This topology agreed with Arshad *et al*. (2009) and Mindell *et al*. (2018), which is unsurprising because in our phylogeny many genera were primarily represented by legacy data generated by Arshad *et al*. (2009), and used by Mindell *et al*. (2018). Notwithstanding, our estimated divergence times were different from those of Mindell *et al*. (2018). For example, we estimated the split between the two main clades of Aegypiinae at 11.7 MYA (vs. 8 MYA, Mindell *et al*., 2018), and the split between *Gyps* and *Necrosyrtes* at 10 MYA (vs. 6 MYA, Mindell *et al*., 2018). Lastly, we estimated that the most recent common ancestor of *Gyps* occurred at 2.7 MYA (vs. 1.5 MYA, Mindell *et al.,* 2018). Arshad *et al*. (2009), who performed multiple divergence estimates using different molecular clock calibrations, arrived at an estimate between 3.7–1.1 MYA.

### AQUILINAE

We recovered a clade containing ten lineages corresponding to genera traditionally classified in the subfamily Aquilinae (*Stephanoaetus*, *Nisaetus*, *Lophotriorchis*, *Polemaetus*, *Spizaetus* [including *S. isidori* (Des Murs, 1845), formerly in the monotypic genus *Oroaetus*; see Haring *et al*., 2007], *Ictinaetus*, *Lophaetus*, *Clanga*, *Aquila* [including *A. africanus* (Cassin, 1865), formerly in *Spizaetus*; see Haring *et al*., 2007], *Hieraaetus*), although generic relationships were different in our phylogeny compared to former studies. Represented only by legacy data, our phylogeny placed the sub-Saharan species Crowned Eagle, *Stephanoaetus coronatus* (Linnaeus, 1766), as sister to *Nisaetus* (Asian hawk-eagles), and this clade was sister to the rest of Aquilinae. This relationship was also found by Helbig *et al*. (2005). However, Haring *et al*. (2007) placed *S. coronatus* as sister to the rest of Aquilinae, excluding the Long-crested Eagle, *Lophaetus occipitalis* (Daudin, 1800); Lerner *et al*. (2017) were unable to consistently determine the placement of *S. coronatus*; and Mindell *et al*. (2018) found *S. coronatus* to be sister to the rest of Aquilinae (including *L. occipitalis*). In our analysis, the Neotropical genus *Spizaetus* was sister to a clade containing all Aquilinae taxa except *S. coronatus* and *Nisaetus*, corroborating several previous studies (Helbig *et al.,* 2005; Lerner *et al.,* 2017; Mindell *et al.,* 2018) but differing from Knapp *et al*. (2019), who relied solely on mitochondrial DNA and recovered *Spizaetus* as sister to *Nisaetus*.

Our 120 taxon UCE phylogeny, which included samples from species representing both major main clades of *Aquila*, and several *Hieraaetus* species, strongly supported (Bootstrap Value = 100) the monophyly of *Aquila*, with *Hieraaetus* as its sister group (similar to Helbig *et al*., 2005). This result differed from Haring *et al*. (2007), Lerner *et al*. (2017), and Mindell *et al*. (2018), who found that *Aquila* was not monophyletic with respect to *Hieraaetus*. Notably, several legacy markers, particularly from the mitochondrial genome, produced topologies that conflicted with the one generated from UCE data, pointing to possible mito-nuclear discord. This discord could explain why existing Aquilinae phylogenies are in conflict. More sequence data are needed to refine our understanding of the relationships within this clade. It is difficult to compare our estimates of divergence timing to former studies with different topologies. However, our estimate for the common ancestor of modern Aquilinae (15.6 MYA) was similar to Knapp *et al*. (2019; *c.*17 MYA), but much older than Mindell *et al*. (2018; *c.*10 MYA).

### HARPIINAE

In our phylogeny, a clade containing the four genera placed in the subfamily Harpiinae (*Harpia Macheiramphus*, *Morphnus*, *Harpyopsis*,) shared a branching order with Griffiths *et al*. (2007) and Mindell *et al*. (2018). However, the placement of Harpiinae within the larger phylogeny differed. We found this clade to be sister to the Buteoninae + Accipitrinae clade, whereas former studies placed it as sister to Aquilinae (Mindell *et al*., 2018) or a clade containing Aquilinae + Buteoninae + Accipitrinae *sensu lato* (Lerner *et al*., 2017) hereafter referring to all members traditionally placed in Melieraxinae, Circinae, and Accipitrinae. Our 120 species UCE phylogeny supported (Bootstrap Value = 100) the placement of Bat Hawk, *Macheiramphus alcinus* Bonaparte, 1850, in the Harpiinae, a relationship previously suggested by several authors on the basis of a single nuclear gene (RAG-1), but with no statistical support (Griffiths *et al*., 2007; Barrowclough *et al*., 2014; Mindell *et al*., 2018). Our divergence estimates for the split between *M. alcinus* and the rest of Harpiinae, and between Crested Eagle, *Morphnus guianensis* (Daudin, 1800), and Harpy Eagle, *Harpia harpyja* (Linnaeus, 1758), —22.1 and 16.4 MYA, respectively—were older than Mindell *et al*.’s (2018) estimates of *c.*15 MYA and *c.*11 MYA, respectively.

### ACCIPITRINAE, CIRCINAE, & MELIERAXINAE

In our 236 species phylogeny, we recovered a clade containing most taxa that were placed within these subfamilies by Lerner & Mindell (2005). In particular, the subfamily Accipitrinae, which included taxa in the genera *Accipiter*, *Erythrotriorchis*, *Megatriorchis*, and *Microspizias*, was rendered polyphyletic by the embedded placement of *Circus* (subfamily Circinae), and several species currently placed within *Accipiter* were not recovered as part of this larger clade. For example, our analysis placed Crested Goshawk, *Accipiter trivirgatus* (Temminck, 1824), as sister to a large clade containing the Accipitrinae (*sensu lato*) and Buteoninae, with high support (Bootstrap Value = 100). This relationship was also recovered, without statistical support, by Mindell *et al*. (2018). Therefore, we join Sangster *et al*. (2021) in applying the generic name *Lophospiza* to [*A.*] *trivirgatus* and Sulawesi Goshawk [*A*.] *griseiceps* (Kaup, 1848), which lacks molecular data but is thought to be closely related based on morphology (Mayr, 1949; Wattel, 1973). We also recommend recognizing these two species in their own subfamily (Lophospizinae).

We recovered a sister relationship between the genera *Harpagus* and *Microspizias*, which was erected by Sangster *et al*. (2021) to accommodate the phylogenetic placement of [*A.*] *superciliosus* and [*A.*] *collaris*. Mindell *et al*. (2018) found a similar relationship, but placed Lizard Buzzard, *Kaupifalco monogrammicus* (Temminck, 1824), within this group. Like Oatley *et al*. (2015), we found the *Microspizias* + *Harpagus* clade to be sister to Buteoninae, whereas Mindell *et al*. (2018) found it to be sister to a clade containing Buteoninae and the rest of Accipitrinae (*sensu lato*). Based on these results, we recognize the genera *Microspizias* and *Harpagus* as members of the subfamily Harpaginae.

Within the clade containing the genera *Melierax*, *Micronisus*, and *Urotriorchis* (all monophyletic), which we classify in the subfamily Melieraxinae, we found that Gabar Goshawk, *Micronisus gabar* (Daudin, 1800), was sister to a clade containing *Melierax* + *Urotriorchis*, confirming a relationship suggested by Mindell *et al*. (2018). However, while Mindell *et al*. (2018) inferred Melieraxinae to be sister to a clade containing Accipitrinae + Circinae + Buteoninae, we found it to be sister to only Accipitrinae + Circinae, an arrangement also proposed by Lerner *et al*. (2008).

In our analysis, *Kaupifalco monogrammicus* was recovered with high support (Bootstrap Value = 100) as sister to a clade containing the remainder of “*Accipiter*” (i.e., excluding [*A.*] *trivirgatus*, [*A*.] *griseiceps*, now in *Lophospiza*, and [*A.*] *superciliosus* and [*A*.] *griseiceps*, now in *Microspizias*) + *Erythrotriorchis* + *Circus* + *Megatriorchis*. This placement conflicted with Griffiths *et al*. (2007) and Mindell *et al*. (2018), who placed *K. monogrammicus* outside the Accipitrinae (*sensu lato*) + Buteoninae clade, and Lerner *et al*. (2008), who placed it as sister to Melieraxinae. Our result was based on UCE data, whereas these former studies used Sanger datasets and found low statistical support for the placement of *K. monogrammicus*.

Even after the removal of the species now placed in *Lophospiza* or *Microspizias*, the remaining members of *Accipiter* (*senso stricto*) still do not form a monophyletic group because *Circus*, *Erythrotriorchis*, and *Megatriorchis* are embedded within the “*Accipiter*” clade. To resolve this issue, we could either (1) synonymize *Circus*, *Erythrotriorchis*, and *Megatriorchis* with *Accipiter*, which would result in the fewest nomenclatural changes; or (2) split *Accipiter* into four genera while retaining *Circus*, *Kaupifalco*, *Erythrotriorchis*, and *Megatriorchis*. In our opinion, the distinct morphological and ecological characteristics of *Circus* necessitate the retention of this genus, despite its nested position within “*Accipiter*”. Additionally, each of these lineages diverged between 15 and 25 MYA, a similar timescale to, or even older than, clades currently treated as genera in other subfamilies. Therefore, we opt for the second approach and propose splitting the remnants of *Accipiter* into multiple genera. This will bring *Accipiter*, which in Clements *et al*. (2021) contained 47 species and was the 11^th^ most speciose genus in the Class Aves, more in line with levels of species-level diversity exhibited by other Accipitridae genera. We provide details about these suggested nomenclatural changes in the following paragraphs.

Within the sister group of *Kaupifalco*, we recovered a clade of “*Accipiter*” species that was sister to a large clade containing the remainder of “*Accipiter*” + *Megatriorchis* + *Circus*. This “*Accipiter*” clade was itself composed of two subclades, which we recognize as sister genera, applying the oldest available generic names for each. One subclade, restricted to sub-Saharan Africa, contained [*A.*] *tachiro* (Daudin, 1800), type species of *Aerospiza* Roberts, 1922, the name we apply to the following species (with new combinations): *Aerospiza tachiro*, *Aerospiza castanilius* (Bonaparte, 1853) **comb. nov.**, *Aerospiza toussenelii* (Verreaux, Verreaux, and Des Murs, 1855) **comb. nov.** The genus *Aerospiza* was erected by Roberts (1922) to include *A. tachiro* and other medium-sized “*Accipiter*” species in Africa, based on the presence of five emarginate primaries, that the fifth primary is longest, that P10 is shorter than the secondaries, and that the tail is ¾ the length of the wing. Wattel (1973) also noted that these species have a long tarsometatarsus and bill, and short middle toe and hallux, relative to other “*Accipiter*” species.

The other subclade of “*Accipiter*” species, which formed part of the sister group of “*Accipiter*” + *Megatriorchis* + *Circus*, included [*A.*] *soloensis* (Horsfield, 1821), type species of the genus *Tachyspiza* Kaup, 1844, which we recognize on the basis of priority, and 26 other species formerly placed in *Accipiter*. Kaup (1844) described two genera, *Tachyspiza* and *Leucospiza* (type species = [*A.*] *novaehollandiae* (Gmelin, JF, 1788), see Sangster *et al*., 2021: 424), of which the type species were placed within this speciose clade. Here, acting as first reviser, we elect to use the name *Tachyspiza*, which is more broadly descriptive of this group of species than *Leucospiza* (i.e., very few members have substantial amounts of white plumage). The genus *Tachyspiza*, as recognized here, is morphologically variable, although members tend to have relatively short toes and talons, especially when compared to *Accipiter* (*sensu stricto*). These two clades (*Aerospiza* and *Tachyspiza*, as denoted here) have been recovered in several previous studies, although the makeup and topology of *Tachyspiza* has varied among studies (Breman *et al*., 2013; Oatley *et al*., 2015; Mindell *et al*., 2018). Our 236 species phylogeny included all three members of *Aerospiza* (one represented by UCEs) and 16 of 27 species now placed in *Tachyspiza* (11 represented by UCEs). Our analysis estimated that the common ancestor of *Aerospiza* and *Tachyspiza* diverged at 17.4 MYA, whereas Mindell *et al*. (2018) gave an estimate of c.10 MYA.

Next, within the sister group of *Erythrotriorchis*, we recovered a clade of “*Accipiter*” species that included *A. nisus*, type species of *Accipiter*, and several other small, primarily bird catching species. We restrict the generic name *Accipiter* to this clade, of which all six species were included in our 236 species phylogeny (three were represented by UCE data). Our topology of *Accipiter* (*sensu stricto*) generally matched Mindell *et al*. (2018), although we found that *A. poliogaster* was a member of this clade rather than sister to *Megatriorchis doriae* Salvadori and D’Albertis, 1876. Notably, this finding is not supported by morphology (Wattel, 1973), but we tentatively place [*A.*] *poliogaster* in *Accipiter* until additional data are available. Similarly, our analysis placed *A. madagascariensis* and *A. ovampensis*, which were represented only by legacy data, within the *Accipiter* (*sensu stricto*) clade, unlike Breman *et al*. (2013), who found them to be sister to a clade containing [*A.*] *gentilis* (without statistical support). In this case, our phylogenetic placement of *A. madagascariensis* and *A. ovampensis* is also supported by morphology, as these two species share the primary characters of *Accipiter* (*sensu stricto*), which are the long, thin tarsometatarsi and toes, relatively small bills and halluces, and small body size.

Within the sister group of *Accipiter* (*sensu stricto*), we recovered a clade of “*Accipiter*” species that was sister to *Megatriorchis* + *Circus*. This topology matched those of Breman *et al*. (2013) and Mindell *et al*. (2018), but had more complete species sampling (7 species, 4 represented by UCEs). For this clade, we apply the generic name *Astur* Lacépède, 1799 (type = *A*. *gentilis*, see Sangster *et al*., 2012: 424) on the basis of priority. As defined, *Astur* is characterized morphologically by their relatively large bills and long halluces. Syringeal characters formerly assumed to be synapomorphic to a clade containing Northern Goshawk (*Astur gentilis*) and Sharp-shinned Hawk (*Accipiter striatus*) are evidently a case of convergence (Griffiths 1994: 794). Within *Astur*, we found two sister clades. The first, which contained *A. gentilis* and its relatives, has a worldwide distribution; the second clade, which contained *A. cooperii* and relatives, is restricted to the Americas. Relative to the “*cooperii* clade”, species in the “*gentilis* clade” are generally distinguished by their larger body size, shorter tarsometatarsi, and shorter middle toes. Although these two clades are genetically and morphologically distinct, we prefer to classify them in one genus because the age of the split (11.6 MYA) is considerably younger than the other genus-level splits in the “*Accipiter*” complex, and younger than most genus-level splits in the family Accipitridae. Should additional research support treating these clades in different genera, the name *Cooperastur* Bonaparte, 1854 is available for the clade containing *A. cooperii* (see Sangster *et al*., 2021: 424).

We found strong support (Bootstrap Value = 100) for a monophyletic clade consisting of all samples in the genus *Circus* de Lacépède, 1799, with comprehensive sampling of extant species and one extinct species. All *Circus* species are associated with open grasslands and/or wetlands, are morphologically united by the presence of a facial disc, and possibly by cranial asymmetry (Pecsics *et al*., 2021). Corroborating Knapp *et al*. (2019), our data indicate that the extinct Eyles’s Harrier, *C. teauteenis* Forbes, 1892, known from New Zealand, was sister of Spotted Harrier, *C. assimilis* Jardine and Selby, 1828, (found in Australia and some islands). In general, our topology agreed with previous studies (Oatley *et al*., 2015; Mindell *et al*., 2018; Knapp *et al*., 2019), with the exception of the placement of *C. assimilis* (and therefore *C. teauteenis*), which we recovered as sister to the rest of *Circus*, rather than sister to one of two main clades within *Circus*. We attribute this difference to more comprehensive sampling of genetic loci in our study. Our divergence estimate for the split between *Circus* and *Megatriorchis* (15.3 MYA), and for the common ancestor of *Circus* (8.7 MYA), were slightly younger than the estimates of Knapp *et al*. (2019; *c.* 17.5 MYA and *c.*10 MYA, respectively), but older than those inferred by Mindell *et al*. (2018; *c.* 10 MYA and *c.*5 MYA respectively) and Oatley *et al*. (2015), who tried several different calibration methods.

Finally, we found that both *Erythrotriorchis* (2 spp.) and *Megatriorchis* (monotypic) were embedded within the “*Accipiter*” (*sensu lato*) + *Circus* clade, as early diverging sisters to larger clades. *Erythrotriorchis* was sister to *Accipiter* (*sensu stricto*) + *Astur* + *Megatriorchis* + *Circus* clade, which conflicted with Mindell *et al*. (2018), who found *Erythrotriorchis* embedded within *Tachyspiza* (as recognized here, see above), and Barrowclough *et al*. (2014), who reconstructed *Erythrotriorchis* as sister to *Tachyspiza*. Both studies lacked statistical support for these relationships. We found that Doria’s Goshawk (*Megatriorchis doriae*) was sister to *Circus* (Bootstrap Value = 100), whereas Mindell *et al*. (2018) placed it as sister to *Accipiter poliogaster*. The placement of *M. doriae* by Mindell *et al*. (2018) was likely an artifact of their supermatrix approach, as these two taxa did not share any sequenced gene regions in their dataset. In contrast, Barrowclough *et al*. (2014) placed *M. doriae* as sister to *Circus*, but did not include any representatives of *Astur*. As expected in studies with radically different tree topologies, divergence estimates published here and in these studies are not easily comparable. We found that the split between *Erythrotriorchis* and its sister group occurred at 20 MYA, whereas *M. doriae* diverged at 15.3 MYA. Conversely, Mindell *et al*. (2018) dated both splits to have occurred more recently than 3.5 MYA.

### BUTEONINAE

Finally, a clade containing all genera traditionally placed in the subfamily Buteoninae was comprised of two major subclades corresponding to the tribes Milvini (*Haliastur*, *Milvus*, *Haliaeetus*) and Buteonini (*Buteo*, *Leucopternis*, *Geranoaetus*, *Pseudastur*, *Parabuteo*, *Rupornis*, *Cryptoleucopteryx, Buteogallus*, *Busarellus*, *Helocolestes*, *Rostrhamus*, *Geranospiza*, *Ictinia*, *Butastur*). With respect to the two tribes and their constituent members, our phylogeny was in agreement with previous studies (Lerner & Mindell, 2005; Amaral *et al*., 2006; Amaral *et al*., 2009; Mindell *et al*., 2018). However, there were some notable differences in the reconstructed relationships within each tribe. For example, Crane Hawk, *Geranospiza caerulescens* (Vieillot, 1817), was sister to a clade containing three monotypic genera (*Busarellus*, *Rostrhamus*, *Helicolestes*) represented by UCE data in our phylogeny. In other studies, *G. caerulescens* was sister to *Buteo* (Riesing *et al*., 2003); or *Rostrhamus* (Lerner & Mindell, 2005; other genera not sampled); or sister to a large clade containing *Busarellus*, *Leucopternis*, *Buteogallus* (*sensu lato*, including two species formerly placed in *Harpyhaliaetus*), *Buteo*, and *Parabuteo* (Amaral *et al*., 2006; other genera not sampled); or in a small clade with *Rostrhamus* and *Busarellus*, that was sister to all Buteonini except *Ictinia* and *Butastur* (Amaral *et al*., 2009); or as sister to *Ictinia*, and with the *Geranospiza* + *Ictinia* clade as sister to the rest of Buteonini except *Butastur* (Mindell *et al*., 2018). Notably, the genus *Helicolestes* was not sampled in any of these former studies.

The enigmatic Barred Hawk, *Morphnarchus princeps* (Sclater, 1865), represented by UCE data in our phylogeny, was sister to a clade containing the monotypic *Cryptoleucopteryx* (also represented by UCE data) and speciose *Buteogallus* (all 9 species represented by legacy data, 6 by UCEs). This relationship was also recovered by Amaral *et al*. (2006), although with no statistical support. Conversely, Amaral *et al*. (2009) placed *M. princeps* as sister to the large clade containing *Buteo*, *Leucopternis*, *Geranoaetus*, *Pseudastur*, *Parabuteo*, and *Rupornis*, with high statistical support. Mindell *et al*. (2018) found the same relationship, but without statistical support. Lastly, the phylogenetic position of Roadside Hawk, *Rupornis magnirostris* (Gmelin, 1788), has been similarly fluid, and remains unresolved. We lacked UCE data for this species (therefore could not assess statistical support), but our analysis of legacy data placed it as sister to a large clade composed of the genera *Buteo*, *Leucopternis*, *Geranoaetus*, *Pseudastur*, and *Parabuteo*. The same arrangement was proposed by Amaral *et al*. (2009). Conversely, Amaral *et al*. (2006) instead found *Parabuteo* to be sister to *Rupornis* + the other genera just mentioned. Mindell *et al*. (2018) placed *Rupornis* as sister to *Parabuteo*, and the *Rupornis* + *Parabuteo* clade as sister to a clade containing *Buteo*, *Leucopternis*, *Geranoaetus*, and *Pseudastur*. Thus, there is unanimous support for the sister relationship of *Buteo* and *Leucopternis*, but the relationships among *Geranoaetus*, *Pseudastur*, and the other genera have been variable. Resolving the relationships of genera in this portion of the phylogeny will require further research, with UCE data from *Rupornis* and *Pseudastur*.

Buteoninae includes several monotypic genera (*Geranospiza*, *Busarellus*, *Rostrhamus*, *Helicolestes*, *Morphnarchus*, *Cryptoleucopteryx*, *Rupornis*) and, at this time, we support retaining all of them. Each of these genera is morphologically, behaviorally, vocally, and ecologically divergent from its closest relatives (del Hoyo *et al*., 1994; Amaral 2009). Furthermore, divergence estimates for the nodes separating these genera (9.3–13 MYA) are older than most generic splits within the Buteoninae (e.g., several pairs of sister genera diverged between 5.6 –8.7 MYA). A notable example is provided by the sister taxa, Slender-billed Kite (*Helicolestes hamatus*) and Snail Kite (*Rostrhamus sociabilis*), which are morphologically and ecologically similar, with diets composed almost exclusively of snails. Whether these genera should be retained, or lumped under the elder name *Rostrhamus* Lesson, 1830, has been a perpetual debate among modern systematists. Recent evaluations by the North and South American Checklist Committees have retained them as monotypic genera (Banks *et al*., 2008: NACC proposal 2007-C-2; Remsen *et al*., 2023: SACC proposal 201). Notably, several committee members indicated they would continue to support the two-genus treatment even if *Helicolestes* and *Rostrhamus* were found to be sister species, unless there was minimal genetic differentiation. Here, with UCE data from both taxa and their closest relatives (i.e., *Busarellus*, *Geranospiza*), we found strong evidence that *Helicolestes* and *Rostrhamus* are sister lineages. Nevertheless, our analysis suggests that they diverged around 10.3 MYA, which is older than some other genus-level splits in Butoninae (see above). Therefore, although *H. hamatus* and *R. sociabilis* share some morphological and ecological characteristics, the preponderance of evidence supports the retention of two, monotypic genera.

## Conclusion

For more than two centuries, scientists have debated the genus-level taxonomy and evolutionary relationships of genera in the diverse family Accipitridae, with little consensus. Here, we reconstructed a strongly-supported phylogeny with UCE data from nearly half of extant species, then using it as a backbone, investigated the phylogenetic relationships of taxa for which legacy (Sanger) sequence data, but no NGS data, were available. We were able to confidently resolve the phylogenetic relationships of 90% of extant species (225 of 249) in Accipitridae, and test the criterion of monophyly for the vast majority of genera. Notably, our UCE sampling included several enigmatic taxa for which little molecular data were previously available and previously had uncertain phylogenetic placements.

The non-monophyly of the diverse genus *Accipiter* is a particularly thorny problem, which has been unresolved for many years (e.g., Griffiths, 2007; Hugall & Stuart-Fox, 2012; Oatley *et al*., 2015; Mindell *et al*., 2018). Heretofore, because of uncertainty caused by limited sampling of taxa and genetic loci, most researchers have deferred taking nomenclatural action to resolve the apparent polyphyly of *Accipiter* (but see Sanger *et al*., 2021). We contend that our combined UCE and legacy datasets are sufficient to resolve this persistent issue, to determine generic boundaries in the polyphyletic *Accipiter* and restrict genus names to monophyletic groups (clades).

Additional sequencing will help further clarify the evolutionary relationships among genera, species, and subspecies in Accipitridae. The taxa in greatest need of additional sampling include the *Accipiter poliogaster*, various monotypic eagle genera, *Pseudastur*, and *Rupornis magnirostris*. Several species within Accipitridae are not monophyletic and a detailed revision of species limits within several clades is sorely needed (e.g., Catanach *et al*., 2021).Divergence time estimates may also be further refined (e.g., by including more fossil calibrations, and identifying datable biogeographic splits).

## Acknowledgements

Funding was provided by NSF grant DEB 2203228, awarded to TAC; and Iridian Genomes grant # IRGEN_RG_2021-1345 Genomic Studies of Eukaryotic Taxa, awarded to SP. TAC was partially supported by NSF grant DEB 1855812, awarded to Jason D. Weckstein. We also thank Sushma Reddy for providing UCE data from *Helicolestes hamatus* and *Kaupifalco monogrammicus*, and Jason Weckstein for providing UCE and/or data from *Accipiter badius*, *A*. *bicolor*, *A. gundlachi*, *A*. *gularis*, *A*. *gundlachi*, *A*. *trivirgatus*, *Circus assimilis*, and *C*. *melanoleucos*. We thank the museums and collection staff that provided samples: Academy of Natural Sciences of Drexel University (Nathan H. Rice), Biodiversity Teaching and Research Collection at Texas A&M University (Gary Voelker and Heather Prestridge), Burke Museum (Sharon Birks), Delaware Museum of Nature & Science (Jean Woods), Field Museum of Natural History (Ben D. Marks, John M. Bates, Shannon J. Hackett), Louisiana State University Museum of Natural Science (Donna Dittmann, Steve Cardiff), Museum of Southwestern Biology (Christopher C. Witt, Andrew B. Johnson), Museum of Vertebrate Zoology (Jeremiah Trimble, Breda M. Zimkus, Emily Blank), Natural History Museum of Los Angeles County (Allison J. Shultz, Kimball L. Garrett), University of Kansas Biodiversity Institute (Mark B. Robbins), Western Foundation of Vertebrate Zoology (Rene Corado), Yale Peabody Museum (Kristof Zyskowski).

## Data Availability

All raw reads generated for this project are deposited in the SRA hosted by NCBI. A list of all accession numbers representing sequences acquired from publicly available sources (ENA, Genbank, and BOLD) is provided in the Supplemental Table 1 within the Legacy_data_samples and UCE_samples_publicly_available tabs.

