## supplemental table 1 for "Enigmas no longer: using Ultraconserved Elements to place several unusual hawk taxa and address the non-monophyly of the genus *Accipiter* (Accipitriformes: Accipitridae)"

### Legacy Data Samples

| sort v2021 | common name | scientific name | RAG-1 | TGFB2 | c-myc | FIB | MB | 12S | 16S | cytb | COI | COII | COIII | ND1 | ND2 | ND3 | ND4 | ND4L | ND5 | ND6 | ATP6 | ATP8 |
|---|---|---|---|---|---|---|---|---|---|---|---|---|---|---|---|---|---|---|---|---|---|---|
| 7409 | California Condor | *Gymnogyps californianus* | SRR14067634 | SRR14067634 | SRR14067634 | SRR14067634 | SRR14067634 | SRR14067634 | SRR14067634 | SRR14067634 | SRR14067634 | SRR14067634 | SRR14067634 | SRR14067634 | SRR14067634 | SRR14067634 | SRR14067634 | SRR14067634 | SRR14067634 | SRR14067634 | SRR14067634 | SRR14067634 |
| 7410 | Andean Condor | *Vultur gryphus* | SRR14067633 | SRR14067633 | SRR14067633 | SRR14067633 | SRR14067633 | SRR14067633 | SRR14067633 | SRR14067633 | SRR14067633 | SRR14067633 | SRR14067633 | SRR14067633 | SRR14067633 | SRR14067633 | SRR14067633 | SRR14067633 | SRR14067633 | SRR14067633 | SRR14067633 | SRR14067633 |
| 7411 | King Vulture | *Sarcoramphus papa* | SRR19167646 | SRR19167646 | SRR19167646 | SRR19167646 | SRR19167646 | SRR19167646 | SRR19167646 | SRR19167646 | SRR19167646 | SRR19167646 | SRR19167646 | SRR19167646 | SRR19167646 | SRR19167646 | SRR19167646 | SRR19167646 | SRR19167646 | SRR19167646 | SRR19167646 | SRR19167646 |
| 7412 | Black Vulture | *Coragyps atratus* | KM876315/DQ881804/SRR8096764 | KX534586 | AY277498/SRR8096764 | EF552759/KX534488/SRR8096764 | SRR8096764 | SRR8096764 | SRR8096764 | SRR8096764 | SRR8096764 | SRR8096764 | SRR8096764 | SRR8096764 | SRR8096764 | SRR8096764 | SRR8096764 | SRR8096764 | SRR8096764 | SRR8096764 | SRR8096764 | SRR8096764 |
| 7416 | Turkey Vulture | *Cathartes aura* | SRR954276 | SRR954276 | SRR954276 | SRR954276 | SRR954276 | SRR954276 | SRR954276 | SRR954276 | SRR954276 | SRR954276 | SRR954276 | SRR954276 | SRR954276 | SRR954276 | SRR954276 | SRR954276 | SRR954276 | SRR954276 | SRR954276 | SRR954276 |
| 7422 | Lesser Yellow-headed Vulture | *Cathartes burrovianus* | | KX534581 | | KX534483 | KX534622 | MN720441 | MN720441 | MN720441 | MN720441 | MN720441 | MN720441 | MN720441 | MN720441 | MN720441 | MN720441 | MN720441 | MN720441 | MN720441 | MN720441 | MN720441 |
| 7425 | Greater Yellow-headed Vulture | *Cathartes melambrotus* | | KX534582 | | KX534484 | KX534623 | | | AF494339/KX534413 | JN801546 | | | | KX534711 | AF076347 | | | | | | |
| 7428 | Secretarybird | *Sagittarius serpentarius* | SRR9946444 | SRR9946444 | SRR9946444 | SRR9946444 | SRR9946444 | SRR9946444 | SRR9946444 | SRR9946444 | SRR9946444 | SRR9946444 | SRR9946444 | SRR9946444 | SRR9946444 | SRR9946444 | SRR9946444 | SRR9946444 | SRR9946444 | SRR9946444 | SRR9946444 | SRR9946444 |
| 7429 | Osprey | *Pandion haliaetus* | SRR17688956 | SRR17688956 | SRR17688956 | SRR17688956 | SRR17688956 | SRR17688956 | SRR17688956 | SRR17688956 | SRR17688956 | SRR17688956 | SRR17688956 | SRR17688956 | SRR17688956 | SRR17688956 | SRR17688956 | SRR17688956 | SRR17688956 | SRR17688956 | SRR17688956 | SRR17688956 |
| 7434 | Pearl Kite | *Gampsonyx swainsonii* | EF078725 | | EU738298 | | | U83723 | | | JN801680/FJ027613 | | | | | | | | | | | |
| 7438 | Black-winged Kite | *Elanus caeruleus* | SRP345212 | SRP345212 | SRP345212 | SRP345212 | SRP345212 | SRP345212 | SRP345212 | SRP345212 | SRP345212 | SRP345212 | SRP345212 | SRP345212 | SRP345212 | SRP345212 | SRP345212 | SRP345212 | SRP345212 | SRP345212 | SRP345212 | SRP345212 |
| 7444 | Black-shouldered Kite | *Elanus axillaris* | SRR18311013 | SRR18311013 | SRR18311013 | SRR18311013 | SRR18311013 | SRR18311013 | SRR18311013 | SRR18311013 | SRR18311013 | SRR18311013 | SRR18311013 | SRR18311013 | SRR18311013 | SRR18311013 | SRR18311013 | SRR18311013 | SRR18311013 | SRR18311013 | SRR18311013 | SRR18311013 |
| 7445 | White-tailed Kite | *Elanus leucurus* | | | GU189422 | AY987163 | | | | AY987233 | FJ027543 | | | | AY987052 | | | | | | | |
| 7448 | Letter-winged Kite | *Elanus scriptus* | | | GU189423 | | | | | | | | | | | | | | | | | |
| 7449 | Scissor-tailed Kite | *Chelictinia riocourii* | MT897865 | | | | | | | MT800533 | MT800541 | | | | | | | | | | | |
| 7450 | African Harrier-Hawk | *Polyboroides typus* | SRR18184967 | SRR18184967 | SRR18184967 | SRR18184967 | SRR18184967 | SRR18184967 | SRR18184967 | SRR18184967 | SRR18184967 | SRR18184967 | SRR18184967 | SRR18184967 | SRR18184967 | SRR18184967 | SRR18184967 | SRR18184967 | SRR18184967 | SRR18184967 | SRR18184967 | SRR18184967 |
| 7454 | Palm-nut Vulture | *Gypohierax angolensis* | EF078729 | | GU189431 | AY987166 | | | | AY987236 | KX012877/KX012878 | | | | AY987055 | | | | | | | |
| 7455 | Madagascar Serpent-Eagle | *Eutriorchis astur* | | | GU189426 | | | | | AY987238 | | | | | AY987057 | | | | | | | |
| 7456 | Bearded Vulture | *Gypaetus barbatus* | EF078728 | | GU189429 | | | | EU144028 | AJ604494 | KX012864 | | | | AY987056 | | | | | | | |
| 7459 | Egyptian Vulture | *Neophron percnopterus* | EF078750 | | GU189430 | AY987165 | | AF173581 | AF173581 | AY987235/EU496468 | KP780411 | | | | AY987054 | | | | | | | |
| 7463 | Hook-billed Kite | *Chondrohierax uncinatus* | | | | AY987167 | | | | EF494393 | | | | | AY987058 | | | | | | | |
| 7467 | Gray-headed Kite | *Leptodon cayanensis* | SRR 17688885 | SRR 17688885 | SRR 17688885 | SRR 17688885 | SRR 17688885 | SRR 17688885 | SRR 17688885 | SRR 17688885 | SRR 17688885 | SRR 17688885 | SRR 17688885 | SRR 17688885 | SRR 17688885 | SRR 17688885 | SRR 17688885 | SRR 17688885 | SRR 17688885 | SRR 17688885 | SRR 17688885 | SRR 17688885 |
| 7471 | European Honey-buzzard | *Pernis apivorus* | EF078753 | | GU189427 | | | U83724 | | X86758 | U83774/GU571536 | | | | AY987061 | | | | | | | |
| 7472 | Sulawesi Honey-buzzard | *Pernis celebensis* | SRR19612714 | SRR19612714 | SRR19612714 | SRR19612714 | SRR19612714 | SRR19612714 | SRR19612714 | SRR19612714 | SRR19612714 | SRR19612714 | SRR19612714 | SRR19612714 | SRR19612714 | SRR19612714 | SRR19612714 | SRR19612714 | SRR19612714 | SRR19612714 | SRR19612714 | SRR19612714 |
| 7473 | Philippine Honey-buzzard | *Pernis steerei* | | | | | | | | AY424390 | | | | | | | | | | | | |
| 7476 | Oriental Honey-buzzard | *Pernis ptilorhynchus* | SRR6650837 | SRR6650837 | SRR6650837 | SRR6650837 | SRR6650837 | SRR6650837 | SRR6650837 | SRR6650837 | SRR6650837 | SRR6650837 | SRR6650837 | SRR6650837 | SRR6650837 | SRR6650837 | SRR6650837 | SRR6650837 | SRR6650837 | SRR6650837 | SRR6650837 | SRR6650837 |
| 7486 | Swallow-tailed Kite | *Elanoides forficatus* | SRR18332084 | SRR18332084 | SRR18332084 | SRR18332084 | SRR18332084 | SRR18332084 | SRR18332084 | SRR18332084 | SRR18332084 | SRR18332084 | SRR18332084 | SRR18332084 | SRR18332084 | SRR18332084 | SRR18332084 | SRR18332084 | SRR18332084 | SRR18332084 | SRR18332084 | SRR18332084 |
| 7489 | Black-breasted Kite | *Hamirostra melanosternon* | EF078734 | | | AY987170 | | | | AY987243 | | | | | AY987062 | | | | | | | |
| 7490 | Long-tailed Honey-buzzard | *Henicopernis longicauda* | SRR18309612 | SRR18309612 | SRR18309612 | SRR18309612 | SRR18309612 | SRR18309612 | SRR18309612 | SRR18309612 | SRR18309612 | SRR18309612 | SRR18309612 | SRR18309612 | SRR18309612 | SRR18309612 | SRR18309612 | SRR18309612 | SRR18309612 | SRR18309612 | SRR18309612 | SRR18309612 |
| 7491 | Black Honey-buzzard | *Henicopernis infuscatus* | KF437487 | | | | | | | | | | | | | | | | | | | |
| 7492 | Square-tailed Kite | *Lophoictinia isura* | EF078744 | | | | | | | AY987244 | | | | | AY987063 | | | | | | | |
| 7493 | African Cuckoo-Hawk | *Aviceda cuculoides* | | | | | | | | AY424398 | | | | | | | | | | | | |
| 7504 | Pacific Baza | *Aviceda subcristata* | SRR19754913 | SRR19754913 | SRR19754913 | SRR19754913 | SRR19754913 | SRR19754913 | SRR19754913 | SRR19754913 | SRR19754913 | SRR19754913 | SRR19754913 | SRR19754913 | SRR19754913 | SRR19754913 | SRR19754913 | SRR19754913 | SRR19754913 | SRR19754913 | SRR19754913 | SRR19754913 |
| 7523 | Red-headed Vulture | *Sarcogyps calvus* | EF078757 | | GU189441 | AY987183 | | | MH423742 | EU496461/AY987264 | KJ506779/MK932903 | | | | AY987083 | | | | | | | |
| 7524 | White-headed Vulture | *Trigonoceps occipitalis* | EF078765 | | GU189443 | | | | | EU496466 | KX012876/KX012846 | | | | AY987084 | | | | | | | |
| 7525 | Cinereous Vulture | *Aegypius monachus* | SRR1251975 | SRR1251975 | SRR1251975 | SRR1251975 | SRR1251975 | SRR1251975 | SRR1251975 | SRR1251975 | SRR1251975 | SRR1251975 | SRR1251975 | SRR1251975 | SRR1251975 | SRR1251975 | SRR1251975 | SRR1251975 | SRR1251975 | SRR1251975 | SRR1251975 | SRR1251975 |
| 7526 | Lappet-faced Vulture | *Torgos tracheliotos* | EF078764 | | GU189442 | AY987185 | | | | AY987267 | HQ168067 | | | | AY987086 | | | | | | | |
| 7530 | Hooded Vulture | *Necrosyrtes monachus* | EF078749 | | GU189440 | AY987178 | | | KJ193294 | EU496459/AY987258 | KX012875/KJ192998 | | | | DQ908990 | | | | | | | |
| 7531 | White-backed Vulture | *Gyps africanus* | EF078730 | | | AY987182 | | | | AY987263 | JQ174984/KX012836 | | | | AY987082 | | | | | | | |
| 7532 | White-rumped Vulture | *Gyps bengalensis* | EU496478 | KX534616 | | AY987179/KX534498 | KX534637 | | | AY987259 | KJ506781/MK932907 | | | | AY987078 | | | | | | | |
| 7533 | Indian Vulture | *Gyps indicus* | EU496477 | | | | | | | DQ908971 | | | | | DQ908989 | | | | | | | |
| 7534 | Slender-billed Vulture | *Gyps tenuirostris* | | | | | | | | DQ908960 | KJ506782 | | | | DQ908982 | | | | | | | |
| 7535 | Rüppells Griffon | *Gyps rueppelli* | SRR17999784 | SRR17999784 | SRR17999784 | SRR17999784 | SRR17999784 | SRR17999784 | SRR17999784 | SRR17999784 | SRR17999784 | SRR17999784 | SRR17999784 | SRR17999784 | SRR17999784 | SRR17999784 | SRR17999784 | SRR17999784 | SRR17999784 | SRR17999784 | SRR17999784 | SRR17999784 |
| 7539 | Himalayan Griffon | *Gyps himalayensis* | EU496476 | | GU189438 | | | KY594709 | KY594709 | KY594709 | KY594709 | KY594709 | KY594709 | KY594709 | KY594709 | KY594709 | KY594709 | KY594709 | KY594709 | KY594709 | KY594709 | KY594709 |
| 7540 | Eurasian Griffon | *Gyps fulvus* | SRR19071310 | SRR19071310 | SRR19071310 | SRR19071310 | SRR19071310 | SRR19071310 | SRR19071310 | SRR19071310 | SRR19071310 | SRR19071310 | SRR19071310 | SRR19071310 | SRR19071310 | SRR19071310 | SRR19071310 | SRR19071310 | SRR19071310 | SRR19071310 | SRR19071310 | SRR19071310 |
| 7545 | Cape Griffon | *Gyps coprotheres* | SRR12539098 | SRR12539098 | SRR12539098 | SRR12539098 | SRR12539098 | SRR12539098 | SRR12539098 | SRR12539098 | SRR12539098 | SRR12539098 | SRR12539098 | SRR12539098 | SRR12539098 | SRR12539098 | SRR12539098 | SRR12539098 | SRR12539098 | SRR12539098 | SRR12539098 | SRR12539098 |
| 7550 | Sulawesi Serpent-Eagle | *Spilornis rufipectus* | | | | | | | | AY987251 | | | | | AY987070 | | | | | | | |
| 7554 | Crested Serpent-Eagle | *Spilornis cheela* | SRR19580237 | SRR19580237 | SRR19580237 | SRR19580237 | SRR19580237 | SRR19580237 | SRR19580237 | SRR19580237 | SRR19580237 | SRR19580237 | SRR19580237 | SRR19580237 | SRR19580237 | SRR19580237 | SRR19580237 | SRR19580237 | SRR19580237 | SRR19580237 | SRR19580237 | SRR19580237 |
| 7577 | Philippine Serpent-Eagle | *Spilornis holospilus* | | | | | | | | AY987249 | HM639911 | | | | AY987068 | | | | | | | |
| 7579 | Philippine Eagle | *Pithecophaga jefferyi* | | | | | | PENDING | PENDING | PENDING | PENDING | PENDING | PENDING | PENDING | PENDING | PENDING | PENDING | PENDING | PENDING | PENDING | PENDING | PENDING |
| 7580 | Bateleur | *Terathopius ecaudatus* | EF078763 | | GU189436 | AY987173 | | | | AY987247 | KX012845/KX012840 | | | | AY987066 | | | | | | | |
| 7581 | Congo Serpent-Eagle | *Dryotriorchis spectabilis* | | | | | | | | AY987255 | | | | | AY987074 | | | | | | | |
| 7584 | Short-toed Snake-Eagle | *Circaetus gallicus* | SRR19071400 | SRR19071400 | SRR19071400 | SRR19071400 | SRR19071400 | SRR19071400 | SRR19071400 | SRR19071400 | SRR19071400 | SRR19071400 | SRR19071400 | SRR19071400 | SRR19071400 | SRR19071400 | SRR19071400 | SRR19071400 | SRR19071400 | SRR19071400 | SRR19071400 | SRR19071400 |
| 7588 | Black-chested Snake-Eagle | *Circaetus pectoralis* | SRR9946917 | SRR9946917 | SRR9946917 | SRR9946917 | SRR9946917 | SRR9946917 | SRR9946917 | SRR9946917 | SRR9946917 | SRR9946917 | SRR9946917 | SRR9946917 | SRR9946917 | SRR9946917 | SRR9946917 | SRR9946917 | SRR9946917 | SRR9946917 | SRR9946917 | SRR9946917 |
| 7589 | Brown Snake-Eagle | *Circaetus cinereus* | | | GU189432 | AY987176 | | | | AY987254 | KX012898 | | | | AY987073 | | | | | | | |
| 7590 | Fasciated Snake-Eagle | *Circaetus fasciolatus* | | | GU189433 | AY987177 | | | | AY987256 | | | | | AY987075 | | | | | | | |
| 7591 | Banded Snake-Eagle | *Circaetus cinerascens* | SRR17913620 | SRR17913620 | SRR17913620 | SRR17913620 | SRR17913620 | SRR17913620 | SRR17913620 | SRR17913620 | SRR17913620 | SRR17913620 | SRR17913620 | SRR17913620 | SRR17913620 | SRR17913620 | SRR17913620 | SRR17913620 | SRR17913620 | SRR17913620 | SRR17913620 | SRR17913620 |
| 7593 | Bat Hawk | *Macheiramphus alcinus* | SRR19077076 | SRR19077076 | SRR19077076 | SRR19077076 | SRR19077076 | SRR19077076 | SRR19077076 | SRR19077076 | SRR19077076 | SRR19077076 | SRR19077076 | SRR19077076 | SRR19077076 | SRR19077076 | SRR19077076 | SRR19077076 | SRR19077076 | SRR19077076 | SRR19077076 | SRR19077076 |
| 7597 | New Guinea Eagle | *Harpyopsis novaeguineae* | EF078737 | SRR17853865 | SRR17853865 | AY987186/SRR17853881 | SRR17853865 | SRR17853867 | SRR17853868 | SRR17853869 | SRR17853870 | SRR17853871 | SRR17853872 | SRR17853873 | SRR17853874 | SRR17853875 | SRR17853876 | SRR17853877 | SRR17853878 | SRR17853879 | SRR17853880 | SRR17853881 |
| 7598 | Crested Eagle | *Morphnus guianensis* | EF078748 | | GU189445 | AY987187 | | | | AJ604496 | | | | | AY987088/AY987089 | | | | | | | |
| 7599 | Harpy Eagle | *Harpia harpyja* | EF078735 | | GU189444 | AY987188 | | | | AJ604495/AY987270 | BOLD IHGTI027-20 | | | | AY987090 | | | | | | | |
| 7600 | Crowned Eagle | *Stephanoaetus coronatus* | EF078762 | | GU189462 | AY987194 | | | | AY987280 | | | | | AY987100 | | | | | | | |
| 7601 | Changeable Hawk-Eagle | *Nisaetus cirrhatus* | SRR17839678 | SRR17839678 | SRR17839678 | SRR17839678 | SRR17839678 | SRR17839678 | SRR17839678 | SRR17839678 | SRR17839678 | SRR17839678 | SRR17839678 | SRR17839678 | SRR17839678 | SRR17839678 | SRR17839678 | SRR17839678 | SRR17839678 | SRR17839678 | SRR17839678 | SRR17839678 |
| 7609 | Flores Hawk-Eagle | *Nisaetus floris* | | | | | | | | AY701118 | | | | | | | | | | | | |
| 7610 | Mountain Hawk-Eagle | *Nisaetus nipalensis* | DRR190839 | DRR190839 | DRR190839 | DRR190839 | DRR190839 | DRR190839 | DRR190839 | DRR190839 | DRR190839 | DRR190839 | DRR190839 | DRR190839 | DRR190839 | DRR190839 | DRR190839 | DRR190839 | DRR190839 | DRR190839 | DRR190839 | DRR190839 |
| 7613 | Legge's Hawk-Eagle | *Nisaetus kelaarti* | | | | | | | | EF459647 | | | | | | | | | | | | |
| 7614 | Blyth's Hawk-Eagle | *Nisaetus alboniger* | | PENDING | | | | PENDING | PENDING | PENDING | PENDING | PENDING | PENDING | PENDING | PENDING | PENDING | PENDING | PENDING | PENDING | PENDING | PENDING | PENDING |
| 7615 | Javan Hawk-Eagle | *Nisaetus bartelsi* | | | | | | | | EF459656 | MT158243 | | | | | | | | | | | |
| 7616 | Sulawesi Hawk-Eagle | *Nisaetus lanceolatus* | | | | | | | | AY987271/AY701130 | | | | | AY987091 | | | | | | | |
| 7617 | Philippine Hawk-Eagle | *Nisaetus philippensis* | AJ601465 | | | | | | | AJ604509 | HM639912 | | | | | | | | | | | |
| 7618 | Pinsker's Hawk-Eagle | *Nisaetus pinskeri* | | | | | | | | AY701127 | | | | | | | | | | | | |
| 7619 | Wallace's Hawk-Eagle | *Nisaetus nanus* | | | | | | | | AY987273 | | | | | AY987093 | | | | | | | |
| 7623 | Martial Eagle | *Polemaetus bellicosus* | EF078754 | | | AY987195 | | | | AJ604504 | | | | | AY987102 | | | | | | | |
| 7624 | Rufous-bellied Eagle | *Lophotriorchis kienerii* | | | | | | | | AY754054 | MK932888 | | | | AY987101 | | | | | | | |
| 7627 | Black Hawk-Eagle | *Spizaetus tyrannus* | SRR9947103 | SRR9947103 | SRR9947103 | SRR9947103 | SRR9947103 | SRR9947103 | SRR9947103 | SRR9947103 | SRR9947103 | SRR9947103 | SRR9947103 | SRR9947103 | SRR9947103 | SRR9947103 | SRR9947103 | SRR9947103 | SRR9947103 | SRR9947103 | SRR9947103 | SRR9947103 |
| 7630 | Ornate Hawk-Eagle | *Spizaetus ornatus* | SRR18186383 | SRR18186383 | SRR18186383 | SRR18186383 | SRR18186383 | SRR18186383 | SRR18186383 | SRR18186383 | SRR18186383 | SRR18186383 | SRR18186383 | SRR18186383 | SRR18186383 | SRR18186383 | SRR18186383 | SRR18186383 | SRR18186383 | SRR18186383 | SRR18186383 | SRR18186383 |
| 7633 | Black-and-white Hawk-Eagle | *Spizaetus melanoleucus* | SRR17913619 | SRR17913619 | SRR17913619 | SRR17913619 | SRR17913619 | SRR17913619 | SRR17913619 | SRR17913619 | SRR17913619 | SRR17913619 | SRR17913619 | SRR17913619 | SRR17913619 | SRR17913619 | SRR17913619 | SRR17913619 | SRR17913619 | SRR17913619 | SRR17913619 | SRR17913619 |
| 7634 | Black-and-chestnut Eagle | *Spizaetus isidori* | EF078751 | | | AY987193 | | | | AJ812238 | BOLD GTICO051-18 | | | | AY987099 | | | | | | | |
| 7636 | Long-crested Eagle | *Lophaetus occipitalis* | EU345542/AJ601458 | | GU189453 | AY987196 | | | | AJ604502 | | | | | AY987103 | | | | | | | |
| 7637 | Black Eagle | *Ictinaetus malaiensis* | EF078739 | | | | | | | EF459611/AY754056 | | | | | AY987109 | | | | | | | |
| 7640 | Lesser Spotted Eagle | *Clanga pomarina* | EU345532/AJ601446 | | GU189452 | | | | | AJ604490 | MK292298 | | | | | | | | | | | |
| 7641 | Indian Spotted Eagle | *Clanga hastata* | | | | | | | | AY987286 | | | | | AY987107 | | | | | | | |
| 7642 | Greater Spotted Eagle | *Clanga clanga* | EU345531/AJ601443 | | | | | | | AJ604487 | MK932912/GU571739 | | | | AY987105 | | | | | | | |
| 7646 | Wahlberg's Eagle | *Hieraaetus wahlbergi* | EU345539/AJ601449 | | GU189458 | AY987202 | | | | AJ604493 | KX012908 | | | | AY987115 | | | | | | | |
| 7647 | Booted Eagle | *Hieraaetus pennatus* | SRR19076291 | SRR19076291 | SRR19076291 | SRR19076291 | SRR19076291 | SRR19076291 | SRR19076291 | SRR19076291 | SRR19076291 | SRR19076291 | SRR19076291 | SRR19076291 | SRR19076291 | SRR19076291 | SRR19076291 | SRR19076291 | SRR19076291 | SRR19076291 | SRR19076291 | SRR19076291 |
| 7648 | Pygmy Eagle | *Hieraaetus weiskei* | | | | | | | | AY987291 | | | | | AY987113 | | | | | | | |
| 7649 | Little Eagle | *Hieraaetus morphnoides* | SRR19076322 | SRR19076322 | SRR19076322 | SRR19076322 | SRR19076322 | SRR19076322 | SRR19076322 | SRR19076322 | SRR19076322 | SRR19076322 | SRR19076322 | SRR19076322 | SRR19076322 | SRR19076322 | SRR19076322 | SRR19076322 | SRR19076322 | SRR19076322 | SRR19076322 | SRR19076322 |
| extinct | Haast's Eagle | *Hieraaetus moorei* | | | | | | MK294166 | MK294166 | MK294166 | MK294166 | MK294166 | MK294166 | MK294166 | MK294166 | MK294166 | MK294166 | MK294166 | MK294166 | MK294166 | MK294166 | MK294166 |
| 7650 | Ayres's Hawk-Eagle | *Hieraaetus ayresii* | SRR19616482 | SRR19616482 | SRR19616482 | SRR19616482 | SRR19616482 | SRR19616482 | SRR19616482 | SRR19616482 | SRR19616482 | SRR19616482 | SRR19616482 | SRR19616482 | SRR19616482 | SRR19616482 | SRR19616482 | SRR19616482 | SRR19616482 | SRR19616482 | SRR19616482 | SRR19616482 |
| 7651 | Tawny Eagle | *Aquila rapax* | EU345527/AJ601447 | | GU189450 | AY987197 | | | | AJ604491/AJ604491 | KX012880 | | | | AY987104 | | | | | MH304920 | | |
| 7655 | Steppe Eagle | *Aquila nipalensis* | SRR18210529 | SRR18210529 | SRR18210529 | SRR18210529 | SRR18210529 | SRR18210529 | SRR18210529 | SRR18210529 | SRR18210529 | SRR18210529 | SRR18210529 | SRR18210529 | SRR18210529 | SRR18210529 | SRR18210529 | SRR18210529 | SRR18210529 | SRR18210529 | SRR18210529 | SRR18210529 |

| | | | | | | | | | | | | | | | | | | | | | | |
|---|---|---|---|---|---|---|---|---|---|---|---|---|---|---|---|---|---|---|---|---|---|---|
| 7659 | Spanish Eagle | *Aquila adalberti* | EU345533 | | | | | | | Z73463 | | | | | | | | | | | | |
| 7660 | Imperial Eagle | *Aquila heliaca* | EU345534/AJ601444 | | GU189448 | AY987198 | | | EU144031 | AJ604488 | KY754483 | | | | AY987106 | | | | | MH304930 | | |
| 7661 | Gurney's Eagle | *Aquila gurneyi* | | | | | | | | AY987303 | | | | | AY987125 | | | | | | | |
| 7662 | Golden Eagle | *Aquila chrysaetos* | ERR3316068 | ERR3316068 | ERR3316068 | ERR3316068 | ERR3316068 | ERR3316068 | ERR3316068 | ERR3316068 | ERR3316068 | ERR3316068 | ERR3316068 | ERR3316068 | ERR3316068 | ERR3316068 | ERR3316068 | ERR3316068 | ERR3316068 | ERR3316068 | ERR3316068 | ERR3316068 |
| 7669 | Wedge-tailed Eagle | *Aquila audax* | SRR19624717 | SRR19624717 | SRR19624717 | SRR19624717 | SRR19624717 | SRR19624717 | SRR19624717 | SRR19624717 | SRR19624717 | SRR19624717 | SRR19624717 | SRR19624717 | SRR19624717 | SRR19624717 | SRR19624717 | SRR19624717 | SRR19624717 | SRR19624717 | SRR19624717 | SRR19624717 |
| 7672 | Cassin's Hawk-Eagle | *Aquila africana* | SRR18186812 | SRR18186812 | SRR18186812 | SRR18186812 | SRR18186812 | SRR18186812 | SRR18186812 | SRR18186812 | SRR18186812 | SRR18186812 | SRR18186812 | SRR18186812 | SRR18186812 | SRR18186812 | SRR18186812 | SRR18186812 | SRR18186812 | SRR18186812 | SRR18186812 | SRR18186812 |
| 7673 | Verreaux's Eagle | *Aquila verreauxii* | EU345525/AJ601448 | | GU189451 | AY987206 | | | | AJ604492 | KX012885 | | | | AY987123 | | | | | | | |
| 7674 | Bonelli's Eagle | *Aquila fasciata* | SRR19580041 | SRR19580041 | SRR19580041 | SRR19580041 | SRR19580041 | SRR19580041 | SRR19580041 | SRR19580041 | SRR19580041 | SRR19580041 | SRR19580041 | SRR19580041 | SRR19580041 | SRR19580041 | SRR19580041 | SRR19580041 | SRR19580041 | SRR19580041 | SRR19580041 | SRR19580041 |
| 7677 | African Hawk-Eagle | *Aquila spilogaster* | SRR18767731 | SRR18767731 | SRR18767731 | SRR18767731 | SRR18767731 | SRR18767731 | SRR18767731 | SRR18767731 | SRR18767731 | SRR18767731 | SRR18767731 | SRR18767731 | SRR18767731 | SRR18767731 | SRR18767731 | SRR18767731 | SRR18767731 | SRR18767731 | SRR18767731 | SRR18767731 |
| 7680 | Lizard Buzzard | *Kaupifalco monogrammicus* | EF078741 | | | | | PENDING | PENDING | PENDING | PENDING | PENDING | PENDING | PENDING | PENDING | PENDING | PENDING | PENDING | PENDING | PENDING | PENDING | PENDING |
| 7683 | Dark Chanting-Goshawk | *Melierax metabates* | SRR19076325 | SRR19076325 | SRR19076325 | SRR19076325 | SRR19076325 | SRR19076325 | SRR19076325 | SRR19076325 | SRR19076325 | SRR19076325 | SRR19076325 | SRR19076325 | SRR19076325 | SRR19076325 | SRR19076325 | SRR19076325 | SRR19076325 | SRR19076325 | SRR19076325 | SRR19076325 |
| 7689 | Eastern Chanting-Goshawk | *Melierax poliopterus* | SRR19077075 | SRR19077075 | SRR19077075 | SRR19077075 | SRR19077075 | SRR19077075 | SRR19077075 | SRR19077075 | SRR19077075 | SRR19077075 | SRR19077075 | SRR19077075 | SRR19077075 | SRR19077075 | SRR19077075 | SRR19077075 | SRR19077075 | SRR19077075 | SRR19077075 | SRR19077075 |
| 7690 | Pale Chanting-Goshawk | *Melierax canorus* | SRR17454566 | | SRR17454566 | SRR17454566 | SRR17454566 | SRR17454566 | SRR17454566 | SRR17454566 | SRR17454566 | SRR17454566 | SRR17454566 | SRR17454566 | SRR17454566 | SRR17454566 | SRR17454566 | SRR17454566 | SRR17454566 | SRR17454566 | SRR17454566 | SRR17454566 |
| 7693 | Gabar Goshawk | *Micronisus gabar* | SRR19076887 | SRR19076887 | SRR19076887 | SRR19076887 | SRR19076887 | SRR19076887 | SRR19076887 | SRR19076887 | SRR19076887 | SRR19076887 | SRR19076887 | SRR19076887 | SRR19076887 | SRR19076887 | SRR19076887 | SRR19076887 | SRR19076887 | SRR19076887 | SRR19076887 | SRR19076887 |
| 7697 | Black-collared Hawk | *Busarellus nigricollis* | SRR18000151 | SRR18000151 | SRR18000151 | SRR18000151 | SRR18000151 | SRR18000151 | SRR18000151 | SRR18000151 | SRR18000151 | SRR18000151 | SRR18000151 | SRR18000151 | SRR18000151 | SRR18000151 | SRR18000151 | SRR18000151 | SRR18000151 | SRR18000151 | SRR18000151 | SRR18000151 |
| 7700 | Snail Kite | *Rostrhamus sociabilis* | SRR18309613 | SRR18309613 | SRR18309613 | SRR18309613 | SRR18309613 | SRR18309613 | SRR18309613 | SRR18309613 | SRR18309613 | SRR18309613 | SRR18309613 | SRR18309613 | SRR18309613 | SRR18309613 | SRR18309613 | SRR18309613 | SRR18309613 | SRR18309613 | SRR18309613 | SRR18309613 |
| 7704 | Slender-billed Kite | *Helicolestes hamatus* | | | | | | PENDING | PENDING | PENDING | PENDING | PENDING | PENDING | PENDING | PENDING | PENDING | PENDING | PENDING | PENDING | PENDING | PENDING | PENDING |
| 7705 | Double-toothed Kite | *Harpagus bidentatus* | SRR18186813 | SRR18186813 | SRR18186813 | SRR18186813 | SRR18186813 | SRR18186813 | SRR18186813 | SRR18186813 | SRR18186813 | SRR18186813 | SRR18186813 | SRR18186813 | SRR18186813 | SRR18186813 | SRR18186813 | SRR18186813 | SRR18186813 | SRR18186813 | SRR18186813 | SRR18186813 |
| 7708 | Rufous-thighed Kite | *Harpagus diodon* | | | | | | | | | | JN801707 | | | | | | | | | | |
| 7709 | Mississippi Kite | *Ictinia mississippiensis* | SRR17688886 | SRR17688886 | SRR17688886 | SRR17688886 | SRR17688886 | SRR17688886 | SRR17688886 | SRR17688886 | SRR17688886 | SRR17688886 | SRR17688886 | SRR17688886 | SRR17688886 | SRR17688886 | SRR17688886 | SRR17688886 | SRR17688886 | SRR17688886 | SRR17688886 | SRR17688886 |
| 7710 | Plumbeous Kite | *Ictinia plumbea* | EF078740 | | GU189506 | AY987219/GQ265061 | | GQ264661 | GQ264661 | GQ264822 | FJ027684/JN801753 | | | | GQ264908 | | | | | GQ264993 | GQ264752 | GQ264752 |
| 7712 | Grasshopper Buzzard | *Butastur rufipennis* | SRR19076324 | SRR19076324 | SRR19076324 | SRR19076324 | SRR19076324 | SRR19076324 | SRR19076324 | SRR19076324 | SRR19076324 | SRR19076324 | SRR19076324 | SRR19076324 | SRR19076324 | SRR19076324 | SRR19076324 | SRR19076324 | SRR19076324 | SRR19076324 | SRR19076324 | SRR19076324 |
| 7713 | White-eyed Buzzard | *Butastur teesa* | EF078716 | | GU189500 | | | | | | EU583328 | | | | EU583267 | | | | | | | |
| 7714 | Rufous-winged Buzzard | *Butastur liventer* | PENDING | PENDING | PENDING | PENDING | PENDING | PENDING | PENDING | PENDING | PENDING | PENDING | PENDING | PENDING | PENDING | PENDING | PENDING | PENDING | PENDING | PENDING | PENDING | PENDING |
| 7715 | Gray-faced Buzzard | *Butastur indicus* | SRR19754896 | SRR19754896 | SRR19754896 | SRR19754896 | SRR19754896 | SRR19754896 | SRR19754896 | SRR19754896 | SRR19754896 | SRR19754896 | SRR19754896 | SRR19754896 | SRR19754896 | SRR19754896 | SRR19754896 | SRR19754896 | SRR19754896 | SRR19754896 | SRR19754896 | SRR19754896 |
| 7716 | Eurasian Marsh-Harrier | *Circus aeruginosus* | EF078722 | KP857800 | GU189481 | KP857746/AY987208 | KP857689 | | U83716 | AY987305 | U83770/GU571336 | | | KP857867 | AY987127 | | | | | | | |
| 7719 | African Marsh-Harrier | *Circus ranivorus* | | KP857817 | | AY987209/KP857765 | KP857708 | | | AY987306 | | | | KP857873 | AY987128 | | | | | | | |
| 7720 | Eastern Marsh-Harrier | *Circus spilonotus* | | | | | KP857734 | LC541452 | LC541452 | LC541452 | LC541452 | LC541452 | LC541452 | LC541452 | LC541452 | LC541452 | LC541452 | LC541452 | LC541452 | LC541452 | LC541452 | LC541452 |
| 7722 | Papuan Marsh-Harrier | *Circus spilothorax* | | KP857818 | | KP857766 | KP857709 | | | | AB921978 | | | KP857875 | | | | | | | | |
| 7724 | Swamp Harrier | *Circus approximans* | SRR19754914 | SRR19754914 | SRR19754914 | SRR19754914 | SRR19754914 | SRR19754914 | SRR19754914 | SRR19754914 | SRR19754914 | SRR19754914 | SRR19754914 | SRR19754914 | SRR19754914 | SRR19754914 | SRR19754914 | SRR19754914 | SRR19754914 | SRR19754914 | SRR19754914 | SRR19754914 |
| 7725 | Reunion Harrier | *Circus maillardi* | SRR21023724 | SRR21023724 | SRR21023724 | SRR21023724 | SRR21023724 | SRR21023724 | SRR21023724 | SRR21023724 | SRR21023724 | SRR21023724 | SRR21023724 | SRR21023724 | SRR21023724 | SRR21023724 | SRR21023724 | SRR21023724 | SRR21023724 | SRR21023724 | SRR21023724 | SRR21023724 |
| 7726 | Malagasy Harrier | *Circus macrosceles* | | KP857840 | | KP857767 | KP857710 | | | | | | | KP857878 | | | | | | | | |
| 7727 | Long-winged Harrier | *Circus buffoni* | | KP857824 | GU189484 | KP857776 | KP857719 | | | | FJ027393 | | | KP857887 | | | | | | | | |
| 7728 | Spotted Harrier | *Circus assimilis* | PENDING | KP857794 | GU189483 | KP857740 | KP857682 | PENDING | PENDING | PENDING | PENDING | PENDING | PENDING | PENDING | PENDING | PENDING | PENDING | PENDING | PENDING | PENDING | PENDING | PENDING |
| extinct | Eyles's Harrier | *Circus teauteensis* | | | | | | MK386464 | MK386464 | MK386464 | MK386464 | MK386464 | MK386464 | MK386464 | MK386464 | MK386464 | MK386464 | MK386464 | MK386464 | MK386464 | MK386464 | MK386464 |
| 7729 | Black Harrier | *Circus maurus* | | KP857809 | | KP857745 | KP857688 | | | | | | | KP857865 | | | | | | | JX878406 | |
| 7730 | Cinereous Harrier | *Circus cinereus* | SRR18191210 | SRR18191210 | SRR18191210 | SRR18191210 | SRR18191210 | SRR18191210 | SRR18191210 | SRR18191210 | SRR18191210 | SRR18191210 | SRR18191210 | SRR18191210 | SRR18191210 | SRR18191210 | SRR18191210 | SRR18191210 | SRR18191210 | SRR18191210 | SRR18191210 | SRR18191210 |
| 7731 | Hen Harrier | *Circus cyaneus* | MT897867 | KP857831 | | KP857782 | KP857701 | KU237286 | KU237286 | KU237286 | KU237286 | KU237286 | KU237286 | KU237286 | KU237286 | KU237286 | KU237286 | KU237286 | KU237286 | KU237286 | KU237286 | KU237286 |
| 7732 | Northern Harrier | *Circus hudsonius* | SRR19170121 | SRR19170121 | SRR19170121 | SRR19170121 | SRR19170121 | SRR19170121 | SRR19170121 | SRR19170121 | SRR19170121 | SRR19170121 | SRR19170121 | SRR19170121 | SRR19170121 | SRR19170121 | SRR19170121 | SRR19170121 | SRR19170121 | SRR19170121 | SRR19170121 | SRR19170121 |
| 7734 | Pallid Harrier | *Circus macrourus* | SRR19076321 | SRR19076321 | SRR19076321 | SRR19076321 | SRR19076321 | SRR19076321 | SRR19076321 | SRR19076321 | SRR19076321 | SRR19076321 | SRR19076321 | SRR19076321 | SRR19076321 | SRR19076321 | SRR19076321 | SRR19076321 | SRR19076321 | SRR19076321 | SRR19076321 | SRR19076321 |
| 7736 | Pied Harrier | *Circus melanoleucos* | | KP857812 | | KP857760 | KP857703 | PENDING | PENDING | PENDING | PENDING | PENDING | PENDING | PENDING | PENDING | PENDING | PENDING | PENDING | PENDING | PENDING | PENDING | PENDING |
| 7737 | Montagu's Harrier | *Circus pygargus* | | KP857829 | GU189487 | KP857780 | KP857704 | | | | JN664079 | | | KP857869 | JN664060 | | | | | | | |
| 7741 | Gray-bellied Hawk | *Accipiter poliogaster* | | | GU189477 | | | | | | JQ173890 | | | | | | | | | | | |
| 7742 | Crested Goshawk | *Accipiter trivirgatus* | | KP857803 | | KP857749 | KP857692 | PENDING | PENDING | PENDING | PENDING | PENDING | PENDING | PENDING | PENDING | PENDING | PENDING | PENDING | PENDING | PENDING | PENDING | PENDING |
| 7755 | Red-chested Goshawk | *Accipiter toussenelii* | EF078710 | KP857822 | GU189480 | KP857773 | KP857717 | | | | JF312130 | | | KP857885 | | | | | | | | |
| 7761 | African Goshawk | *Accipiter tachiro* | | KP857816 | | KP857764 | KP857707 | | | | JF312199 | | | KP857872 | | | | | | | | |
| 7768 | Chestnut-flanked Sparrowhawk | *Accipiter castanilius* | SRR18769175 | SRR18769175 | SRR18769175 | SRR18769175 | SRR18769175 | SRR18769175 | SRR18769175 | SRR18769175 | SRR18769175 | SRR18769175 | SRR18769175 | SRR18769175 | SRR18769175 | SRR18769175 | SRR18769175 | SRR18769175 | SRR18769175 | SRR18769175 | SRR18769175 | SRR18769175 |
| 7769 | Shikra | *Accipiter badius* | PENDING | KP857836 | GU189467 | KP857787 | KP857731 | PENDING | PENDING | PENDING | PENDING | PENDING | PENDING | PENDING | PENDING | PENDING | PENDING | PENDING | PENDING | PENDING | PENDING | PENDING |
| 7781 | Levant Sparrowhawk | *Accipiter brevipes* | | | GU189469 | | | PENDING | PENDING | PENDING | PENDING | PENDING | PENDING | PENDING | PENDING | PENDING | PENDING | PENDING | PENDING | PENDING | PENDING | PENDING |
| 7782 | Chinese Sparrowhawk | *Accipiter soloensis* | SRR6650832 | SRR6650832 | SRR6650832 | SRR6650832 | SRR6650832 | SRR6650832 | SRR6650832 | SRR6650832 | SRR6650832 | SRR6650832 | SRR6650832 | SRR6650832 | SRR6650832 | SRR6650832 | SRR6650832 | SRR6650832 | SRR6650832 | SRR6650832 | SRR6650832 | SRR6650832 |
| 7783 | Frances's Sparrowhawk | *Accipiter francesiae* | | | GU189471 | | | | | | JF792346 | | | | | | | | | | | |
| 7789 | Variable Goshawk | *Accipiter hiogaster* | SRR18310035 | SRR18310035 | SRR18310035 | SRR18310035 | SRR18310035 | SRR18310035 | SRR18310035 | SRR18310035 | SRR18310035 | SRR18310035 | SRR18310035 | SRR18310035 | SRR18310035 | SRR18310035 | SRR18310035 | SRR18310035 | SRR18310035 | SRR18310035 | SRR18310035 | SRR18310035 |
| 7813 | Gray Goshawk | *Accipiter novaehollandiae* | SRR17913812 | SRR17913812 | SRR17913812 | SRR17913812 | SRR17913812 | SRR17913812 | SRR17913812 | SRR17913812 | SRR17913812 | SRR17913812 | SRR17913812 | SRR17913812 | SRR17913812 | SRR17913812 | SRR17913812 | SRR17913812 | SRR17913812 | SRR17913812 | SRR17913812 | SRR17913812 |
| 7814 | Brown Goshawk | *Accipiter fasciatus* | SRR19076323 | SRR19076323 | SRR19076323 | SRR19076323 | SRR19076323 | SRR19076323 | SRR19076323 | SRR19076323 | SRR19076323 | SRR19076323 | SRR19076323 | SRR19076323 | SRR19076323 | SRR19076323 | SRR19076323 | SRR19076323 | SRR19076323 | SRR19076323 | SRR19076323 | SRR19076323 |
| 7827 | Black-mantled Goshawk | *Accipiter melanochlamys* | SRR18000157 | SRR18000157 | SRR18000157 | SRR18000157 | SRR18000157 | SRR18000157 | SRR18000157 | SRR18000157 | SRR18000157 | SRR18000157 | SRR18000157 | SRR18000157 | SRR18000157 | SRR18000157 | SRR18000157 | SRR18000157 | SRR18000157 | SRR18000157 | SRR18000157 | SRR18000157 |
| 7836 | New Caledonia Goshawk | *Accipiter haplochrous* | | KP857827 | | KP857778 | KP857722 | | | | | | | KP857890 | | | | | | | | |
| 7837 | Fiji Goshawk | *Accipiter rufitorques* | SRR18191211 | SRR18191211 | SRR18191211 | SRR18191211 | SRR18191211 | SRR18191211 | SRR18191211 | SRR18191211 | SRR18191211 | SRR18191211 | SRR18191211 | SRR18191211 | SRR18191211 | SRR18191211 | SRR18191211 | SRR18191211 | SRR18191211 | SRR18191211 | SRR18191211 | SRR18191211 |
| 7840 | Imitator Sparrowhawk | *Accipiter imitator* | EF078707 | | | | | | | | | | | | | | | | | | | |
| 7841 | Gray-headed Goshawk | *Accipiter poliocephalus* | | KP857802 | | KP857748 | KP857691 | | | | | | | KP857856 | | | | | | | | |
| 7843 | Tiny Hawk | *Accipiter superciliosus* | SRR20912048 | SRR20912048 | SRR20912048 | SRR20912048 | SRR20912048 | SRR20912048 | SRR20912048 | SRR20912048 | SRR20912048 | SRR20912048 | SRR20912048 | SRR20912048 | SRR20912048 | SRR20912048 | SRR20912048 | SRR20912048 | SRR20912048 | SRR20912048 | SRR20912048 | SRR20912048 |
| 7846 | Semicollared Hawk | *Accipiter collaris* | | | | | | | | | BOLD IAVHB205-14 | | | | | | | | | | | |
| 7847 | Red-thighed Sparrowhawk | *Accipiter erythropus* | | KP857833 | | KP857784 | KP857728 | | | | JF312107 | | | KP857896 | | | | | | | | |
| 7850 | Little Sparrowhawk | *Accipiter minullus* | | KP857839 | | KP857751 | KP857694 | | | | JF312158 | | | KP857859 | | | | | | | | |
| 7851 | Japanese Sparrowhawk | *Accipiter gularis* | | | | | | PENDING | PENDING | PENDING | PENDING | PENDING | PENDING | PENDING | PENDING | PENDING | PENDING | PENDING | PENDING | PENDING | PENDING | PENDING |
| 7856 | Besra | *Accipiter virgatus* | SRR21034869 | SRR21034869 | SRR21034869 | SRR21034869 | SRR21034869 | SRR21034869 | SRR21034869 | SRR21034869 | SRR21034869 | SRR21034869 | SRR21034869 | SRR21034869 | SRR21034869 | SRR21034869 | SRR21034869 | SRR21034869 | SRR21034869 | SRR21034869 | SRR21034869 | SRR21034869 |
| 7883 | Madagascar Sparrowhawk | *Accipiter madagascariensis* | | | | | | | | | JF792339 | | | | | | | | | | | |
| 7884 | Ovambo Sparrowhawk | *Accipiter ovampensis* | | | GU189476 | | | | | | JF312121 | | | | | | | | | | | |
| 7885 | Eurasian Sparrowhawk | *Accipiter nisus* | DRR190838 | DRR190838 | DRR190838 | DRR190838 | DRR190838 | DRR190838 | DRR190838 | DRR190838 | DRR190838 | DRR190838 | DRR190838 | DRR190838 | DRR190838 | DRR190838 | DRR190838 | DRR190838 | DRR190838 | DRR190838 | DRR190838 | DRR190838 |
| 7893 | Rufous-breasted Sparrowhawk | *Accipiter rufiventris* | SRR17688967 | SRR17688967 | SRR17688967 | SRR17688967 | SRR17688967 | SRR17688967 | SRR17688967 | SRR17688967 | SRR17688967 | SRR17688967 | SRR17688967 | SRR17688967 | SRR17688967 | SRR17688967 | SRR17688967 | SRR17688967 | SRR17688967 | SRR17688967 | SRR17688967 | SRR17688967 |
| 7896 | Sharp-shinned Hawk | *Accipiter striatus* | SRR17839736 | SRR17839736 | SRR17839736 | SRR17839736 | SRR17839736 | SRR17839736 | SRR17839736 | SRR17839736 | SRR17839736 | SRR17839736 | SRR17839736 | SRR17839736 | SRR17839736 | SRR17839736 | SRR17839736 | SRR17839736 | SRR17839736 | SRR17839736 | SRR17839736 | SRR17839736 |
| 7909 | Cooper's Hawk | *Accipiter cooperii* | SRR17454565 | SRR17454565 | SRR17454565 | SRR17454565 | SRR17454565 | SRR17454565 | SRR17454565 | SRR17454565 | SRR17454565 | SRR17454565 | SRR17454565 | SRR17454565 | SRR17454565 | SRR17454565 | SRR17454565 | SRR17454565 | SRR17454565 | SRR17454565 | SRR17454565 | SRR17454565 |
| 7911 | Gundlach's Hawk | *Accipiter gundlachi* | | | | | | PENDING | PENDING | PENDING | PENDING | PENDING | PENDING | PENDING | PENDING | PENDING | PENDING | PENDING | PENDING | PENDING | PENDING | PENDING |
| 7914 | Bicolored Hawk | *Accipiter bicolor* | | KP857821 | GU189468 | KP857772/AY987210 | KP857716 | PENDING | PENDING | PENDING | PENDING | PENDING | PENDING | PENDING | PENDING | PENDING | PENDING | PENDING | PENDING | PENDING | PENDING | PENDING |
| 7922 | Black Goshawk | *Accipiter melanoleucus* | SRR16687839 | SRR16687839 | SRR16687839 | SRR16687839 | SRR16687839 | SRR16687839 | SRR16687839 | SRR16687839 | SRR16687839 | SRR16687839 | SRR16687839 | SRR16687839 | SRR16687839 | SRR16687839 | SRR16687839 | SRR16687839 | SRR16687839 | SRR16687839 | SRR16687839 | SRR16687839 |
| 7925 | Henst's Goshawk | *Accipiter henstii* | | | GU189473 | | | | | MK433177 | | | | | | | | | | | | |
| 7926 | Northern Goshawk | *Accipiter gentilis* | DRR191147 | DRR191147 | DRR191147 | DRR191147 | DRR191147 | DRR191147 | DRR191147 | DRR191147 | DRR191147 | DRR191147 | DRR191147 | DRR191147 | DRR191147 | DRR191147 | DRR191147 | DRR191147 | DRR191147 | DRR191147 | DRR191147 | DRR191147 |
| 7939 | Meyer's Goshawk | *Accipiter meyerianus* | | | | | | | | MK433189 | | | | | | | | | | | | |
| 7941 | Chestnut-shouldered Goshawk | *Erythrotriorchis buergersi* | SRR22494351 | SRR22494351 | SRR22494351 | SRR22494351 | SRR22494351 | SRR22494351 | SRR22494351 | SRR22494351 | SRR22494351 | SRR22494351 | SRR22494351 | SRR22494351 | SRR22494351 | SRR22494351 | SRR22494351 | SRR22494351 | SRR22494351 | SRR22494351 | SRR22494351 | SRR22494351 |
| 7942 | Red Goshawk | *Erythrotriorchis radiatus* | KF437489 | | | | | | | | | | | | | | | | | | | |
| 7943 | Doria's Goshawk | *Megatriorchis doriae* | SRR19070526 | SRR19070526 | SRR19070526 | SRR19070526 | SRR19070526 | SRR19070526 | SRR19070526 | SRR19070526 | SRR19070526 | SRR19070526 | SRR19070526 | SRR19070526 | SRR19070526 | SRR19070526 | SRR19070526 | SRR19070526 | SRR19070526 | SRR19070526 | SRR19070526 | SRR19070526 |
| 7944 | Long-tailed Hawk | *Urotriorchis macrourus* | | | | | | | | EU583379 | | | | | EU583318 | | | | | | | |
| 7945 | Red Kite | *Milvus milvus* | SRR18000158 | SRR18000158 | SRR18000158 | SRR18000158 | SRR18000158 | SRR18000158 | SRR18000158 | SRR18000158 | SRR18000158 | SRR18000158 | SRR18000158 | SRR18000158 | SRR18000158 | SRR18000158 | SRR18000158 | SRR18000158 | SRR18000158 | SRR18000158 | SRR18000158 | SRR18000158 |
| 7948 | Black Kite | *Milvus migrans* | SRR6650833 | SRR6650833 | SRR6650833 | SRR6650833 | SRR6650833 | SRR6650833 | SRR6650833 | SRR6650833 | SRR6650833 | SRR6650833 | SRR6650833 | SRR6650833 | SRR6650833 | SRR6650833 | SRR6650833 | SRR6650833 | SRR6650833 | SRR6650833 | SRR6650833 | SRR6650833 |
| 7960 | Whistling Kite | *Haliastur sphenurus* | SRR17835005 | SRR17835005 | SRR17835005 | SRR17835005 | SRR17835005 | SRR17835005 | SRR17835005 | SRR17835005 | SRR17835005 | SRR17835005 | SRR17835005 | SRR17835005 | SRR17835005 | SRR17835005 | SRR17835005 | SRR17835005 | SRR17835005 | SRR17835005 | SRR17835005 | SRR17835005 |
| 7961 | Brahminy Kite | *Haliastur indus* | SRR17688966 | SRR17688966 | SRR17688966 | SRR17688966 | SRR17688966 | SRR17688966 | SRR17688966 | SRR17688966 | SRR17688966 | SRR17688966 | SRR17688966 | SRR17688966 | SRR17688966 | SRR17688966 | SRR17688966 | SRR17688966 | SRR17688966 | SRR17688966 | SRR17688966 | SRR17688966 |
| 7967 | Bald Eagle | *Haliaeetus leucocephalus* | SRR1176806 | SRR1176806 | SRR1176806 | SRR1176806 | SRR1176806 | SRR1176806 | SRR1176806 | SRR1176806 | SRR1176806 | SRR1176806 | SRR1176806 | SRR1176806 | SRR1176806 | SRR1176806 | SRR1176806 | SRR1176806 | SRR1176806 | SRR1176806 | SRR1176806 | SRR1176806 |
| 7971 | White-tailed Eagle | *Haliaeetus albicilla* | SRR6650836 | SRR6650836 | SRR6650836 | SRR6650836 | SRR6650836 | SRR6650836 | SRR6650836 | SRR6650836 | SRR6650836 | SRR6650836 | SRR6650836 | SRR6650836 | SRR6650836 | SRR6650836 | SRR6650836 | SRR6650836 | SRR6650836 | SRR6650836 | SRR6650836 | SRR6650836 |
| 7972 | Pallas's Fish-Eagle | *Haliaeetus leucoryphus* | | | | | | | | Z73469 | | | | | AY987135 | | | | | | | |
| 7973 | Steller's Sea-Eagle | *Haliaeetus pelagicus* | DRR250432 | DRR250433 | DRR250434 | DRR250435 | DRR250436 | DRR250438 | DRR250439 | DRR250440 | DRR250441 | DRR250442 | DRR250443 | DRR250444 | DRR250445 | DRR250446 | DRR250447 | DRR250448 | DRR250449 | DRR250450 | DRR250451 | DRR250452 |
| 7974 | White-bellied Sea-Eagle | *Haliaeetus leucogaster* | SRR19076271 | SRR19076272 | SRR19076273 | SRR19076274 | SRR19076275 | SRR19076277 | SRR19076278 | SRR19076279 | SRR19076280 | SRR19076281 | SRR19076282 | SRR19076283 | SRR19076284 | SRR19076285 | SRR19076286 | SRR19076287 | SRR19076288 | SRR19076289 | SRR19076290 | SRR19076291 |
| 7975 | Sanford's Sea-Eagle | *Haliaeetus sanfordi* | | | | | | | | AY987322 | | | | | AY987144 | | | | | | | |
| 7976 | African Fish-Eagle | *Haliaeetus vocifer* | ERR2688507 | ERR2688507 | ERR2688507 | ERR2688507 | ERR2688507 | ERR2688507 | ERR2688507 | ERR2688507 | ERR2688507 | ERR2688507 | ERR2688507 | ERR2688507 | ERR2688507 | ERR2688507 | ERR2688507 | ERR2688507 | ERR2688507 | ERR2688507 | ERR2688507 | ERR2688507 |
| 7977 | Madagascar Fish-Eagle | *Haliaeetus vociferoides* | | | GU189493 | AY987217 | | | | AY987320 | | | | | AY987142 | | | | | | | |

| | | | | | | | | | | | | | | | | | | | | | | |
|---|---|---|---|---|---|---|---|---|---|---|---|---|---|---|---|---|---|---|---|---|---|---|
| 7978 | Lesser Fish-Eagle | *Haliaeetus humilis* | | | | | | | | AY987317 | | | | | AY987139 | | | | | | | |
| 7981 | Gray-headed Fish-Eagle | *Haliaeetus ichthyaetus* | SRR19625954 | SRR19625954 | SRR19625954 | SRR19625954 | SRR19625954 | SRR19625954 | SRR19625954 | SRR19625954 | SRR19625954 | SRR19625954 | SRR19625954 | SRR19625954 | SRR19625954 | SRR19625954 | SRR19625954 | SRR19625954 | SRR19625954 | SRR19625954 | SRR19625954 | SRR19625954 |
| 7984 | Crane Hawk | *Geranospiza caerulescens* | SRR18000150 | SRR18000150 | SRR18000150 | SRR18000150 | SRR18000150 | SRR18000150 | SRR18000150 | SRR18000150 | SRR18000150 | SRR18000150 | SRR18000150 | SRR18000150 | SRR18000150 | SRR18000150 | SRR18000150 | SRR18000150 | SRR18000150 | SRR18000150 | SRR18000150 | SRR18000150 |
| 7993 | Plumbeous Hawk | *Cryptoleucopteryx plumbea* | SRR19616483 | SRR19616483 | SRR19616483 | SRR19616483 | SRR19616483 | SRR19616483 | SRR19616483 | SRR19616483 | SRR19616483 | SRR19616483 | SRR19616483 | SRR19616483 | SRR19616483 | SRR19616483 | SRR19616483 | SRR19616483 | SRR19616483 | SRR19616483 | SRR19616483 | SRR19616483 |
| 7994 | Slate-colored Hawk | *Buteogallus schistaceus* | | | | GQ265077/EU583254 | | DQ148361/GQ264685 | GQ264685 | EU583372 | | | | | EU583311/GQ264931 | | | | | DQ148331 | DQ148390 | DQ148390 |
| 7995 | Common Black Hawk | *Buteogallus anthracinus* | EF078720 | | GU189496 | GQ265019/AY987223 | | GQ264605 | GQ264605 | AY987327/GQ264778 | DQ432785 | | | | AY987149/GQ264863 | | | | | GQ264948 | GQ264703 | GQ264703 |
| 8003 | Cuban Black Hawk | *Buteogallus gundlachii* | SRR21034923 | SRR21034923 | SRR21034923 | SRR21034923 | SRR21034923 | SRR21034923 | SRR21034923 | SRR21034923 | SRR21034923 | SRR21034923 | SRR21034923 | SRR21034923 | SRR21034923 | SRR21034923 | SRR21034923 | SRR21034923 | SRR21034923 | SRR21034923 | SRR21034923 | SRR21034923 |
| 8004 | Rufous Crab Hawk | *Buteogallus aequinoctialis* | SRR18184966 | SRR18184966 | SRR18184966 | SRR18184966 | SRR18184966 | SRR18184966 | SRR18184966 | SRR18184966 | SRR18184966 | SRR18184966 | SRR18184966 | SRR18184966 | SRR18184966 | SRR18184966 | SRR18184966 | SRR18184966 | SRR18184966 | SRR18184966 | SRR18184966 | SRR18184966 |
| 8005 | Savanna Hawk | *Buteogallus meridionalis* | SRR21034868 | SRR21034868 | SRR21034868 | SRR21034868 | SRR21034868 | SRR21034868 | SRR21034868 | SRR21034868 | SRR21034868 | SRR21034868 | SRR21034868 | SRR21034868 | SRR21034868 | SRR21034868 | SRR21034868 | SRR21034868 | SRR21034868 | SRR21034868 | SRR21034868 | SRR21034868 |
| 8006 | White-necked Hawk | *Buteogallus lacernulatus* | SRR18186811 | SRR18186811 | SRR18186811 | SRR18186811 | SRR18186811 | SRR18186811 | SRR18186811 | SRR18186811 | SRR18186811 | SRR18186811 | SRR18186811 | SRR18186811 | SRR18186811 | SRR18186811 | SRR18186811 | SRR18186811 | SRR18186811 | SRR18186811 | SRR18186811 | SRR18186811 |
| 8007 | Great Black Hawk | *Buteogallus urubitinga* | | | GU189497 | AY987222/GQ265048 | | GQ264647 | GQ264647 | GQ264808 | FJ027240 | | | | GQ264894 | | | | | GQ264984 | GQ264743 | GQ264743 |
| 8011 | Solitary Eagle | *Buteogallus solitarius* | SRR17835758 | SRR17835758 | SRR17835758 | SRR17835758 | SRR17835758 | SRR17835758 | SRR17835758 | SRR17835758 | SRR17835758 | SRR17835758 | SRR17835758 | SRR17835758 | SRR17835758 | SRR17835758 | SRR17835758 | SRR17835758 | SRR17835758 | SRR17835758 | SRR17835758 | SRR17835758 |
| 8014 | Chaco Eagle | *Buteogallus coronatus* | SRR17853877 | SRR17853877 | SRR17853877 | SRR17853877 | SRR17853877 | SRR17853877 | SRR17853877 | SRR17853877 | SRR17853877 | SRR17853877 | SRR17853877 | SRR17853877 | SRR17853877 | SRR17853877 | SRR17853877 | SRR17853877 | SRR17853877 | SRR17853877 | SRR17853877 | SRR17853877 |
| 8016 | Barred Hawk | *Morphnarchus princeps* | SRR21034931 | SRR21034931 | SRR21034931 | SRR21034931 | SRR21034931 | SRR21034931 | SRR21034931 | SRR21034931 | SRR21034931 | SRR21034931 | SRR21034931 | SRR21034931 | SRR21034931 | SRR21034931 | SRR21034931 | SRR21034931 | SRR21034931 | SRR21034931 | SRR21034931 | SRR21034931 |
| 8017 | Roadside Hawk | *Rupornis magnirostris* | EF078719 | | | GQ265031/AY987225 | | DQ148352/GQ264626 | GQ264626 | GQ264790 | FJ027248/JN801525 | | | | GQ264877 | | | | | DQ148322 | DQ148382/GQ264723 | DQ148382/GQ264723 |
| 8032 | Harris's Hawk | *Parabuteo unicinctus* | SRR18333723 | SRR18333723 | SRR18333723 | SRR18333723 | SRR18333723 | SRR18333723 | SRR18333723 | SRR18333723 | SRR18333723 | SRR18333723 | SRR18333723 | SRR18333723 | SRR18333723 | SRR18333723 | SRR18333723 | SRR18333723 | SRR18333723 | SRR18333723 | SRR18333723 | SRR18333723 |
| 8035 | White-rumped Hawk | *Parabuteo leucorrhous* | | | | GQ265080 | | GQ264690 | GQ264690 | GQ264850 | FJ027243 | | | | GQ264936 | | | | | GQ265008 | GQ264767 | GQ264767 |
| 8036 | White-tailed Hawk | *Geranoaetus albicaudatus* | SRR18332296 | SRR18332296 | SRR18332296 | SRR18332296 | SRR18332296 | SRR18332296 | SRR18332296 | SRR18332296 | SRR18332296 | SRR18332296 | SRR18332296 | SRR18332296 | SRR18332296 | SRR18332296 | SRR18332296 | SRR18332296 | SRR18332296 | SRR18332296 | SRR18332296 | SRR18332296 |
| 8040 | Variable Hawk | *Geranoaetus polyosoma* | SRR19170547 | SRR19170547 | SRR19170547 | SRR19170547 | SRR19170547 | SRR19170547 | SRR19170547 | SRR19170547 | SRR19170547 | SRR19170547 | SRR19170547 | SRR19170547 | SRR19170547 | SRR19170547 | SRR19170547 | SRR19170547 | SRR19170547 | SRR19170547 | SRR19170547 | SRR19170547 |
| 8045 | Black-chested Buzzard-Eagle | *Geranoaetus melanoleucus* | EF078726 | | | GQ265053/AY987226 | | GQ264652 | GQ264652 | GQ264813 | FJ027626 | | | | GQ264899 | | | | | GQ264986 | GQ264745 | GQ264745 |
| 8049 | Mantled Hawk | *Pseudastur polionotus* | | | | EU583252/GQ265075 | | DQ148348/GQ264682 | GQ264682 | GQ264842 | | | | | GQ264928 | | | | | DQ148318 | DQ148376 | DQ148376 |
| 8050 | White Hawk | *Pseudastur albicollis* | | | GU189507 | GQ265064/AY987227 | | GQ264662 | GQ264662 | GQ264823 | JN801773 | | | | GQ264909 | | | | | GQ264994 | GQ264753 | GQ264753 |
| 8055 | Gray-backed Hawk | *Pseudastur occidentalis* | SRR21115777 | SRR21115777 | SRR21115777 | EU583247/GQ265073 | SRR21115777 | GQ264678 | GQ264678 | EU583363/GQ264839 | | | | | EU583302/GQ264925 | | | | | GQ265002 | GQ264761 | GQ264761 |
| 8056 | Semiplumbeous Hawk | *Leucopternis semiplumbeus* | | | | GQ265078/EU583255 | | DQ148359/GQ264687 | GQ264687 | GQ264847 | JQ175261 | | | | GQ264933 | | | | | DQ148329 | DQ148388/DQ148389 | DQ148388/DQ148389 |
| 8057 | Black-faced Hawk | *Leucopternis melanops* | SRR18272076 | SRR18272076 | SRR18272076 | SRR18272076 | SRR18272076 | SRR18272076 | SRR18272076 | SRR18272076 | SRR18272076 | SRR18272076 | SRR18272076 | SRR18272076 | SRR18272076 | SRR18272076 | SRR18272076 | SRR18272076 | SRR18272076 | SRR18272076 | SRR18272076 | SRR18272076 |
| 8058 | White-browed Hawk | *Leucopternis kuhli* | SRR18272075 | SRR18272075 | SRR18272075 | SRR18272075 | SRR18272075 | SRR18272075 | SRR18272075 | SRR18272075 | SRR18272075 | SRR18272075 | SRR18272075 | SRR18272075 | SRR18272075 | SRR18272075 | SRR18272075 | SRR18272075 | SRR18272075 | SRR18272075 | SRR18272075 | SRR18272075 |
| 8060 | Gray Hawk | *Buteo plagiatus* | | | | | | | | | DQ433339 | | | | | | | | | | | |
| 8061 | Gray-lined Hawk | *Buteo nitidus* | SRR17853888 | SRR17853888 | SRR17853888 | SRR17853888 | SRR17853888 | SRR17853888 | SRR17853888 | SRR17853888 | SRR17853888 | SRR17853888 | SRR17853888 | SRR17853888 | SRR17853888 | SRR17853888 | SRR17853888 | SRR17853888 | SRR17853888 | SRR17853888 | SRR17853888 | SRR17853888 |
| 8066 | Red-shouldered Hawk | *Buteo lineatus* | SRR19616268 | SRR19616268 | SRR19616268 | SRR19616268 | SRR19616268 | SRR19616268 | SRR19616268 | SRR19616268 | SRR19616268 | SRR19616268 | SRR19616268 | SRR19616268 | SRR19616268 | SRR19616268 | SRR19616268 | SRR19616268 | SRR19616268 | SRR19616268 | SRR19616268 | SRR19616268 |
| 8074 | Ridgway's Hawk | *Buteo ridgwayi* | | | | GQ265041 | | GQ264636 | GQ264636 | GQ264801 | | | | | GQ264887 | | | | | GQ264975 | GQ264732 | GQ264732 |
| 8075 | Broad-winged Hawk | *Buteo platypterus* | SRR18186351 | SRR18186351 | SRR18186351 | SRR18186351 | SRR18186351 | SRR18186351 | SRR18186351 | SRR18186351 | SRR18186351 | SRR18186351 | SRR18186351 | SRR18186351 | SRR18186351 | SRR18186351 | SRR18186351 | SRR18186351 | SRR18186351 | SRR18186351 | SRR18186351 | SRR18186351 |
| 8085 | Hawaiian Hawk | *Buteo solitarius* | | | | | | GQ264643 | GQ264643 | GQ264804 | JQ174219 | | | | GQ264890 | | | | | GQ264980 | GQ264739 | GQ264739 |
| 8086 | Short-tailed Hawk | *Buteo brachyurus* | | | | GQ265022 | | GQ264609 | GQ264609 | GQ264782 | DQ432783 | | | | GQ264867 | | | | | GQ264952 | GQ264707 | GQ264707 |
| 8089 | White-throated Hawk | *Buteo albigula* | SRR18000144 | SRR18000144 | SRR18000144 | SRR18000144 | SRR18000144 | SRR18000144 | SRR18000144 | SRR18000144 | SRR18000144 | SRR18000144 | SRR18000144 | SRR18000144 | SRR18000144 | SRR18000144 | SRR18000144 | SRR18000144 | SRR18000144 | SRR18000144 | SRR18000144 | SRR18000144 |
| 8090 | Swainson's Hawk | *Buteo swainsoni* | SRR18726455 | SRR18726455 | SRR18726455 | SRR18726455 | SRR18726455 | SRR18726455 | SRR18726455 | SRR18726455 | SRR18726455 | SRR18726455 | SRR18726455 | SRR18726455 | SRR18726455 | SRR18726455 | SRR18726455 | SRR18726455 | SRR18726455 | SRR18726455 | SRR18726455 | SRR18726455 |
| 8091 | Galapagos Hawk | *Buteo galapagoensis* | | | | GQ265023 | | GQ264613 | GQ264613 | GQ264783 | AY870866/AY870870 | | | | GQ264868 | | | | | GQ264956 | GQ264711 | GQ264711 |
| 8092 | Zone-tailed Hawk | *Buteo albonotatus* | SRR18332297 | SRR18332297 | SRR18332297 | SRR18332297 | SRR18332297 | SRR18332297 | SRR18332297 | SRR18332297 | SRR18332297 | SRR18332297 | SRR18332297 | SRR18332297 | SRR18332297 | SRR18332297 | SRR18332297 | SRR18332297 | SRR18332297 | SRR18332297 | SRR18332297 | SRR18332297 |
| 8093 | Rufous-tailed Hawk | *Buteo ventralis* | SRR17454564 | SRR17454564 | SRR17454564 | SRR17454564 | SRR17454564 | SRR17454564 | SRR17454564 | SRR17454564 | SRR17454564 | SRR17454564 | SRR17454564 | SRR17454564 | SRR17454564 | SRR17454564 | SRR17454564 | SRR17454564 | SRR17454564 | SRR17454564 | SRR17454564 | SRR17454564 |
| 8094 | Red-tailed Hawk | *Buteo jamaicensis* | SRR17913811 | SRR17913811 | SRR17913811 | SRR17913811 | SRR17913811 | SRR17913811 | SRR17913811 | SRR17913811 | SRR17913811 | SRR17913811 | SRR17913811 | SRR17913811 | SRR17913811 | SRR17913811 | SRR17913811 | SRR17913811 | SRR17913811 | SRR17913811 | SRR17913811 | SRR17913811 |
| 8114 | Rough-legged Hawk | *Buteo lagopus* | SRR21034922 | SRR21034922 | SRR21034922 | SRR21034922 | SRR21034922 | SRR21034922 | SRR21034922 | SRR21034922 | SRR21034922 | SRR21034922 | SRR21034922 | SRR21034922 | SRR21034922 | SRR21034922 | SRR21034922 | SRR21034922 | SRR21034922 | SRR21034922 | SRR21034922 | SRR21034922 |
| 8121 | Ferruginous Hawk | *Buteo regalis* | SRR19167630 | SRR19167630 | SRR19167630 | SRR19167630 | SRR19167630 | SRR19167630 | SRR19167630 | SRR19167630 | SRR19167630 | SRR19167630 | SRR19167630 | SRR19167630 | SRR19167630 | SRR19167630 | SRR19167630 | SRR19167630 | SRR19167630 | SRR19167630 | SRR19167630 | SRR19167630 |
| 8123 | Common Buzzard | *Buteo buteo* | SRR19071506 | SRR19071506 | SRR19071506 | SRR19071506 | SRR19071506 | SRR19071506 | SRR19071506 | SRR19071506 | SRR19071506 | SRR19071506 | SRR19071506 | SRR19071506 | SRR19071506 | SRR19071506 | SRR19071506 | SRR19071506 | SRR19071506 | SRR19071506 | SRR19071506 | SRR19071506 |
| 8131 | Himalayan Buzzard | *Buteo refectus* | | | | | | KM364882 | KM364882 | KM364882 | KM364882 | KM364882 | KM364882 | KM364882 | KM364882 | KM364882 | KM364882 | KM364882 | KM364882 | KM364882 | KM364882 | KM364882 |
| 8133 | Eastern Buzzard | *Buteo japonicus* | SRR6656033 | SRR6656033 | SRR6656033 | SRR6656033 | SRR6656033 | SRR6656033 | SRR6656033 | SRR6656033 | SRR6656033 | SRR6656033 | SRR6656033 | SRR6656033 | SRR6656033 | SRR6656033 | SRR6656033 | SRR6656033 | SRR6656033 | SRR6656033 | SRR6656033 | SRR6656033 |
| 8139 | Mountain Buzzard | *Buteo oreophilus* | SRR19067422 | SRR19067422 | SRR19067422 | SRR19067422 | SRR19067422 | SRR19067422 | SRR19067422 | SRR19067422 | SRR19067422 | SRR19067422 | SRR19067422 | SRR19067422 | SRR19067422 | SRR19067422 | SRR19067422 | SRR19067422 | SRR19067422 | SRR19067422 | SRR19067422 | SRR19067422 |
| 8140 | Forest Buzzard | *Buteo trizonatus* | | | | EU583231 | | | | EU583339 | | | | | EU583278 | | | | | | | |
| 8145 | Long-legged Buzzard | *Buteo rufinus* | SRR19067741 | SRR19067741 | SRR19067741 | SRR19067741 | SRR19067741 | SRR19067741 | SRR19067741 | SRR19067741 | SRR19067741 | SRR19067741 | SRR19067741 | SRR19067741 | SRR19067741 | SRR19067741 | SRR19067741 | SRR19067741 | SRR19067741 | SRR19067741 | SRR19067741 | SRR19067741 |
| 8150 | Upland Buzzard | *Buteo hemilasius* | SRR19067439 | SRR19067439 | SRR19067439 | SRR19067439 | SRR19067439 | SRR19067439 | SRR19067439 | SRR19067439 | SRR19067439 | SRR19067439 | SRR19067439 | SRR19067439 | SRR19067439 | SRR19067439 | SRR19067439 | SRR19067439 | SRR19067439 | SRR19067439 | SRR19067439 | SRR19067439 |
| 8153 | Red-necked Buzzard | *Buteo auguralis* | | | | | | | | | | | | | | | | | | AY213016 | | |
| 8154 | Augur Buzzard | *Buteo augur* | | | | GQ265021 | | GQ264607 | GQ264607 | GQ264780 | | | | | GQ264865 | | | | | GQ264950 | GQ264705 | GQ264705 |
| 8155 | Archer's Buzzard | *Buteo archeri* | | | | | | | | | | | | | | | | | | AY216930 | | |
| 8156 | Jackal Buzzard | *Buteo rufofuscus* | | | | EU583237 | | GQ264639 | | EU583342 | | | | | EU583281 | | | | | GQ264978 | GQ264735 | GQ264735 |

### Legacy data mined from NGS

| sort v2021 | common name | scientific name | Voucher | Locality | Type | ENA accession |
|---|---|---|---|---|---|---|
| 7579 | Philippine Eagle | *Pithecophaga jefferyi* | ANSP 174095 | Philippines: Cebu | Toepad | PENDING |
| 7614 | Blyth's Hawk-Eagle | *Nisaetus alboniger* | WFVZ 42302 | Malaysia: Sabah | Toepad | PENDING |
| 7911 | Gundlach's Hawk | *Accipiter gundlachi* | LSU 141446 | Cuba: Villa Clara | Toepad | PENDING |
| 7914 | Bicolored Hawk | *Accipiter bicolor* | LSU 24224 | Mexico: Oaxaca | Toepad | PENDING |
| 7714 | Rufous-winged Buzzard | *Butastur liventer* | ANSP 127461 | Thailand: Chanuman | Toepad | PENDING |

### UCE samples this study

| sort v2021 | common name | scientific name | Voucher | Locality | Type | ENA accession |
|---|---|---|---|---|---|---|
| 7411 | King Vulture | *Sarcoramphus papa* | MSB: Bird: 27868 | Peru: San Martin | Tissue | SRR 19167646 |
| 7429 | Osprey | *Pandion haliaetus* | ANSP 205240 | USA: Maryland | Tissue | SRR 17688956 |
| 7444 | Black-shouldered Kite | *Elanus axillaris* | KU Birds 98282 | AUSTRALIA: New South Wales | Tissue | SRR 18311013 |
| 7450 | African Harrier-Hawk | *Polyboroides typus* | ANSP 23822 | Equatorial Guinea: Centro Sur | Tissue | SRR 18184967 |
| 7467 | Gray-headed Kite | *Leptodon cayanensis* | ANSP 187788 | Guyana: Potaro-Siparuni | Tissue | SRR 17688885 |
| 7472 | Sulawesi Honey-buzzard | *Pernis celebensis* | MCZ 270340 | Indonesia: Central Sulawesi | Toepad | SRR 19612714 |
| 7486 | Swallow-tailed Kite | *Elanoides forficatus* | TCWC Birds 21022 | USA: Texas | Tissue | SRR 18332084 |
| 7490 | Long-tailed Honey-buzzard | *Henicopernis longicauda* | KU Birds 96967 | Papua New Guinea: Oro | Tissue | SRR 18309612 |
| 7504 | Pacific Baza | *Aviceda subcristata* | KU Birds 98819 | AUSTRALIA: New South Wales | Tissue | SRR 19754913 |
| 7535 | Rüppells Griffon | *Gyps rueppelli* | LACM Birds 61059 | Kenya: Rift Valley | Toepad | SRR 17999784 |
| 7540 | Eurasian Griffon | *Gyps fulvus* | KU Birds 91202 | Spain: Madrid | Tissue | SRR 19071310 |
| 7554 | Crested Serpent-Eagle | *Spilornis cheela* | MCZ 178153 | Indonesia: Aceh | Toepad | SRR 19580237 |
| 7584 | Short-toed Snake-Eagle | *Circaetus gallicus* | TCWC 29791 | Italy | Tissue | SRR 19071400 |
| 7591 | Banded Snake-Eagle | *Circaetus cinerascens* | LACM Birds 64143 | Uganda: Western | Toepad | SRR 17913620 |
| 7593 | Bat Hawk | *Macheiramphus alcinus* | WFVZ 37194 | Malaysia: Sabah | Toepad | SRR 19077076 |
| 7597 | New Guinea Eagle | *Harpyopsis novaeguineae* | LACM Birds TISSUE 2025 | PAPUA NEW GUINEA: WEST SEPIK | Tissue | SRR 17853881 |
| 7601 | Changeable Hawk-Eagle | *Nisaetus cirrhatus* | LACM Birds 32809 | India: MADHYA PRADESH | Toepad | SRR 17839678 |
| 7630 | Ornate Hawk-Eagle | *Spizaetus ornatus* | ANSP 188268 | Guyana: Potaro-Siparuni | Tissue | SRR 18186383 |
| 7633 | Black-and-white Hawk-Eagle | *Spizaetus melanoleucus* | ANSP 187773 | Guyana: Potaro-Siparuni | Tissue | SRR 17913619 |
| 7647 | Booted Eagle | *Hieraaetus pennatus* | KU Birds 91934 | Spain: Madrid | Tissue | SRR 19076291 |
| 7649 | Little Eagle | *Hieraaetus morphnoides* | KU Birds 99982 | AUSTRALIA: New South Wales | Tissue | SRR 19076322 |
| 7650 | Ayres's Hawk-Eagle | *Hieraaetus ayresii* | LACM Birds 54485 | Kenya: Rift Valley | Toepad | SRR 19616482 |
| 7655 | Steppe Eagle | *Aquila nipalensis* | KU Birds 119895 | Mongolia: Gobi Altai | Tissue | SRR 18210529 |
| 7669 | Wedge-tailed Eagle | *Aquila audax* | ANSP 192576 | Australia: Tasmania | Tissue | SRR 19624717 |
| 7672 | Cassin's Hawk-Eagle | *Aquila africana* | LACM Birds 72636 | Uganda: Western | Toepad | SRR 18186812 |
| 7674 | Bonelli's Eagle | *Aquila fasciata* | MCZ 261829 | Indonesia: East Nusa Tenggara | Toepad | SRR 19580041 |
| 7677 | African Hawk-Eagle | *Aquila spilogaster* | LACM Birds 75152 | Kenya: Rift Valley | Tissue | SRR 18767731 |
| 7680 | Lizard Buzzard | *Kaupifalco monogrammicus* | FMNH 452458 | Malawi: Chitipa | Tissue | PENDING |
| 7683 | Dark Chanting-Goshawk | *Melierax metabates* | KU Birds 70660 | Kenya: Kongelai | Toepad | SRR 19076325 |
| 7689 | Eastern Chanting-Goshawk | *Melierax poliopterus* | LACM Birds 88623 | Kenya: Eastern | Toepad | SRR 19077075 |
| 7690 | Pale Chanting-Goshawk | *Melierax canorus* | LACM Birds 79106 | Namibia: Karas | Toepad | SRR17454566 |
| 7693 | Gabar Goshawk | *Micronisus gabar* | KU Birds 110795 | Ghana: Upper West | Tissue | SRR 19076887 |

| | | | | | | |
|---|---|---|---|---|---|---|
| 7697 | Black-collared Hawk | *Busarellus nigricollis* | KU Birds 91625 | Paraguay: Alto Paraguay | Tissue | SRR 18000151 |
| 7700 | Snail Kite | *Rostrhamus sociabilis* | KU Birds 90149 | Paraguay: Ñeembucú | Tissue | SRR 18309613 |
| 7704 | Slender-billed Kite | *Helicolestes hamatus* | FMNH 100848 | Brazil: Amazonas | Toepad | PENDING |
| 7705 | Double-toothed Kite | *Harpagus bidentatus* | ANSP 31945 | Brazil: Amazonas | Toepad | SRR 18186813 |
| 7709 | Mississippi Kite | *Ictinia mississippiensis* | ANSP 205149 | USA: Texas | Tissue | SRR 17688886 |
| 7712 | Grasshopper Buzzard | *Butastur rufipennis* | KU Birds 110797 | Ghana: Upper West | Tissue | SRR 19076324 |
| 7715 | Gray-faced Buzzard | *Butastur indicus* | KU Birds 98340 | Philippines: Negros Island | Tissue | SRR 19754896 |
| 7724 | Swamp Harrier | *Circus approximans* | KU Birds 114996 | New Zealand: West Coast | Tissue | SRR 19754914 |
| 7725 | Reunion Harrier | *Circus maillardi* | ANSP 133 | Réunion | Toepad | SRR 21023724 |
| 7728 | Spotted Harrier | *Circus assimilis* | ANSP 189865 | Australia, South Australia | Tissue | PENDING |
| 7730 | Cinereous Harrier | *Circus cinereus* | KU Birds 78213 | Argentina: Chubut | Toepad | SRR 18191210 |
| 7732 | Northern Harrier | *Circus hudsonius* | ANSP 206397 | USA: Minnesota | Tissue | SRR 19170121 |
| 7734 | Pallid Harrier | *Circus macrourus* | LACM 41658 | Chad: Ennedi-Ouest | Toepad | SRR 19076321 |
| 7736 | Pied Harrier | *Circus melanoleucos* | UWBM 75277 | Russia: Primorskiy Kray | Tissue | PENDING |
| 7742 | Crested Goshawk | *Accipiter trivirgatus* | DMNH 14455 | Philippines: Quezon | Toepad | PENDING |
| 7768 | Chestnut-flanked Sparrowhawk | *Accipiter castanilius* | ANSP 190275 | Equatorial Guinea: Centro Sur | Tissue | SRR 18769175 |
| 7769 | Shikra | *Accipiter badius* | DMNH 57032 | Nigeria: Oyo | Toepad | PENDING |
| 7781 | Levant Sparrowhawk | *Accipiter brevipes* | YPM 141224 | Russia: Rostovskaya Oblast | Tissue | PENDING |
| 7789 | Variable Goshawk | *Accipiter hiogaster* | KU Birds 97031 | Papua New Guinea: Oro | Tissue | SRR 18310035 |
| 7813 | Gray Goshawk | *Accipiter novaehollandiae* | ANSP 192569 | Australia: Tasmania | Tissue | SRR 17913812 |
| 7814 | Brown Goshawk | *Accipiter fasciatus* | ANSP 27428 | Australia: Queensland | Tissue | SRR 19076323 |
| 7827 | Black-mantled Goshawk | *Accipiter melanochlamys* | LACM Birds 78235 | Kenya:Eastern | Toepad | SRR 18000157 |
| 7837 | Fiji Goshawk | *Accipiter rufitorques* | KU Birds 120222 | Fiji: Vanua Levu | Tissue | SRR 18191211 |
| 7843 | Tiny Hawk | *Accipiter superciliosus* | ANSP 35662 | Brazil: Maranhao | | SRR 20912048 |
| 7851 | Japanese Sparrowhawk | *Accipiter gularis* | UWBM 59854 | MONGOLIA: Dornod Aymag | Tissue | PENDING |
| 7856 | Besra | *Accipiter virgatus* | KU Birds 122241 | Philippines: Mindanao | Tissue | SRR 21034869 |
| 7893 | Rufous-breasted Sparrowhawk | *Accipiter rufiventris* | LACM Birds 75162 | Kenya: Rift Valley | Toepad | SRR 17688967 |
| 7896 | Sharp-shinned Hawk | *Accipiter striatus* | LSU 171160 | Bolivia: Santa Cruz | Tissue | SRR 17839736 |
| 7909 | Cooper's Hawk | *Accipiter cooperii* | ANSP 204237 | USA: Washington | Tissue | SRR 17454565 |
| 7914 | Bicolored Hawk | *Accipiter bicolor* | FMNH 260973 | Colombia: Norte de Santander | Toepad | PENDING |
| 7922 | Black Goshawk | *Accipiter melanoleucus* | KU Birds 125095 | Papua New Guinea: East Highlands | Tissue | SRR 16687839 |
| 7941 | Chestnut-shouldered Goshawk | *Erythrotriorchis buergersi* | BBM-NG-54096 | Papua New Guinea: Morobe | Toepad | SRR 22494351 |
| 7943 | Doria's Goshawk | *Megatriorchis doriae* | KU Birds 96989 | Papua New Guinea: Oro | Tissue | SRR 19070526 |
| 7945 | Red Kite | *Milvus milvus* | KU Birds 91933 | Spain: Madrid | Tissue | SRR 18000158 |

| | | | | | | |
|---|---|---|---|---|---|---|
| 7960 | Whistling Kite | *Haliastur sphenurus* | ANSP 191939 | AUSTRALIA: New South Wales | Tissue | SRR 17835005 |
| 7961 | Brahminy Kite | *Haliastur indus* | ANSP 191935 | Australia: Queensland | Tissue | SRR 17688966 |
| 7974 | White-bellied Sea-Eagle | *Haliaeetus leucogaster* | KU Birds 110339 | AUSTRALIA: New South Wales | Tissue | SRR 19076292 |
| 7981 | Gray-headed Fish-Eagle | *Haliaeetus ichthyaetus* | WFVZ 37202 | Malaysia: Sabah | Tissue | SRR 19625954 |
| 7984 | Crane Hawk | *Geranospiza caerulescens* | KU Birds 94799 | Guyana: Barima-Waini | Tissue | SRR 18000150 |
| 7993 | Plumbeous Hawk | *Cryptoleucopteryx plumbea* | ANSP 182230 | Ecuador: Esmeraldas | Toepad | SRR 19616483 |
| 8003 | Cuban Black Hawk | *Buteogallus gundlachii* | ANSP 160276 | Cuba: Isla de la Juventud | Toepad | SRR 21034923 |
| 8004 | Rufous Crab Hawk | *Buteogallus aequinoctialis* | ANSP 187793 | Guyana: Mahaica-Berbice | Tissue | SRR 18184966 |
| 8005 | Savanna Hawk | *Buteogallus meridionalis* | ANSP 34402 | Brazil: Para | Tissue | SRR 21034868 |
| 8006 | White-necked Hawk | *Buteogallus lacernulatus* | LACM Birds 27764 | Brazil: Rio de Janeiro | Toepad | SRR18186811 |
| 8011 | Solitary Eagle | *Buteogallus solitarius* | LSU B22903 | Bolivia: La Paz | Tissue | SRR 17835758 |
| 8014 | Chaco Eagle | *Buteogallus coronatus* | LACM Birds 114879 | Captive | Tissue | SRR 17853877 |
| 8016 | Barred Hawk | *Morphnarchus princeps* | ANSP 180156 | Ecuador: Esmeraldas | Toepad | SRR 21034931 |
| 8032 | Harris's Hawk | *Parabuteo unicinctus* | TCWC Birds 14879 | USA: Texas | Tissue | SRR 18333723 |
| 8036 | White-tailed Hawk | *Geranoaetus albicaudatus* | TCWC Birds 16353 | USA: Texas | Tissue | SRR 18332296 |
| 8040 | Variable Hawk | *Geranoaetus polyosoma* | MSB:Bird:35942 | Peru: Lima | Tissue | SRR 19170547 |
| 8057 | Black-faced Hawk | *Leucopternis melanops* | ANSP 187658 | Guyana: Potaro-Siparuni | Tissue | SRR 18272076 |
| 8058 | White-browed Hawk | *Leucopternis kuhli* | ANSP 206505 | Brazil: Amazonas | Tissue | SRR 18272075 |
| 8061 | Gray-lined Hawk | *Buteo nitidus* | ANSP 188341 | Guyana: Potaro-Siparuni | Tissue | SRR 17853888 |
| 8066 | Red-shouldered Hawk | *Buteo lineatus* | ANSP 206271 | USA: Pennsylvania | Tissue | SRR 19616268 |
| 8075 | Broad-winged Hawk | *Buteo platypterus* | ANSP 193958 | USA: New Jersey | Tissue | SRR 18186351 |
| 8089 | White-throated Hawk | *Buteo albigula* | KU Birds 127901 | Peru: Ayacucho | Tissue | SRR 18000144 |
| 8090 | Swainson's Hawk | *Buteo swainsoni* | ANSP 192342 | USA: New Mexico | Tissue | SRR 18726455 |
| 8092 | Zone-tailed Hawk | *Buteo albonotatus* | TCWC Birds 15864 | USA: Texas | Tissue | SRR 18332297 |
| 8093 | Rufous-tailed Hawk | *Buteo ventralis* | LACM Birds 56760 | Argentina: Río Negro | Toepad | SRR 17454564 |
| 8094 | Red-tailed Hawk | *Buteo jamaicensis* | ANSP 195233 | USA: Pennsylvania | Tissue | SRR 17913811 |
| 8114 | Rough-legged Hawk | *Buteo lagopus* | ANSP 206395 | USA: North Dakota | Tissue | SRR 21034922 |
| 8121 | Ferruginous Hawk | *Buteo regalis* | MSB:Bird:39066 | USA: New Mexico | Tissue | SRR 19167630 |
| 8123 | Common Buzzard | *Buteo buteo* | TCWC 29790 | Italy | Tissue | SRR 19071506 |
| 8139 | Mountain Buzzard | *Buteo oreophilus* | LACM Birds 75171 | Kenya: Rift Valley | Toepad | SRR 19067422 |
| 8145 | Long-legged Buzzard | *Buteo rufinus* | KU Birds 119971 | Mongolia: Gobi Altai | Tissue | SRR 19067741 |
| 8150 | Upland Buzzard | *Buteo hemilasius* | KU Birds 120830 | Mongolia: Bayanhangor | Tissue | SRR 19067439 |

### UCE samples publicly available

| sort v2021 | common name | scientific name | ENA accession |
|---|---|---|---|
| 7409 | California Condor | *Gymnogyps californianus* | SRR 14067634 |
| 7410 | Andean Condor | *Vultur gryphus* | SRR 14067633 |
| 7412 | Black Vulture | *Coragyps atratus* | SRR 8096764 |
| 7416 | Turkey Vulture | *Cathartes aura* | SRR 954276 |
| 7428 | Secretarybird | *Sagittarius serpentarius* | SRR 9946444 |
| 7438 | Black-winged Kite | *Elanus caeruleus* | SRP 345212 |
| 7476 | Oriental Honey-buzzard | *Pernis ptilorhynchus* | SRR 6650837 |
| 7525 | Cinereous Vulture | *Aegypius monachus* | SRR 1251975 |
| 7545 | Cape Griffon | *Gyps coprotheres* | SRR 12539098 |
| 7588 | Black-chested Snake-Eagle | *Circaetus pectoralis* | SRR 9946917 |
| 7610 | Mountain Hawk-Eagle | *Nisaetus nipalensis* | DRR 190839 |
| 7627 | Black Hawk-Eagle | *Spizaetus tyrannus* | SRR 9947103 |
| 7662 | Golden Eagle | *Aquila chrysaetos* | ERR 3316068 |
| 7782 | Chinese Sparrowhawk | *Accipiter soloensis* | SRR 6650832 |
| 7885 | Eurasian Sparrowhawk | *Accipiter nisus* | DRR 190838 |
| 7926 | Northern Goshawk | *Accipiter gentilis* | DRR 191147 |
| 7948 | Black Kite | *Milvus migrans* | SRR 6650833 |
| 7967 | Bald Eagle | *Haliaeetus leucocephalus* | SRR 1176806 |
| 7971 | White-tailed Eagle | *Haliaeetus albicilla* | SRR 6650836 |
| 7973 | Steller's Sea-Eagle | *Haliaeetus pelagicus* | DRR 250453 |
| 7976 | African Fish-Eagle | *Haliaeetus vocifer* | ERR 2688507 |
| 8133 | Eastern Buzzard | *Buteo japonicus* | SRR 6656033 |
